# A conserved photosynthetic cytochrome enhances growth of *Chlamydomonas reinhardtii* in fluctuating light

**DOI:** 10.64898/2026.02.13.705717

**Authors:** Darius Kosmützky, Laura T Wey, Lauri Nikkanen, Aron Ferenczi, Joshua M Lawrence, Alberto Scarampi, Attila Molnar, Yagut Allahverdiyeva, Christopher J Howe

## Abstract

Having at least one of the small soluble electron carriers cytochrome *c*_6_ (*c*_6_) and plastocyanin is indispensable for photosynthesis. It was believed that *c*_6_ had been lost from plants until a ubiquitous and highly conserved homologue, now named cytochrome *c*_6A_ (*c*_6A_), was found in *Arabidopsis thaliana*. In spite of an early suggestion that *c*_6A_ functioned like *c*_6_ in the photosynthetic electron transport chain it was soon shown to be unable to do so despite the strong structural homology between the two proteins. The function of *c*_6A_, and why it is apparently universal in plants and green algae, therefore remained unknown. Here, we show that, in the green alga *Chlamydomonas reinhardtii*, *c*_6A_ confers a growth advantage under fluctuating light and is crucial for maintaining the light harvesting balance between photosystems I and II in photomixotrophic conditions. We show that in the absence of *c*_6A_, the light harvesting balance shifts towards PSII, leading to a more reduced plastoquinone pool and increased photooxidative stress. This study provides insights into how photosynthetic organisms acclimatize to stressful light conditions, indicating that *c*_6A_ is important for this adaptation. These findings provide a basis for further mechanistic studies on a hypothesized role for *c*_6A_ in thiol-based redox regulation in the thylakoid lumen, with implications for photoprotection mechanisms such as state transitions.

## Introduction

The photosynthetic electron transport chain (PETC) is essential for nearly all life on Earth (Fig. 1a). At least one of the soluble electron carriers plastocyanin and cytochrome *c*_6_ is indispensable for the PETC to function (1, 2). A close homologue of *c*_6_, cytochrome *c*_6A_, was first discovered in *Arabidopsis thaliana* and later in other plants and eukaryotic algae (3, 4). Whereas plants lost the gene for *c*_6_ and contain only *c*_6A_, some eukaryotic algae such as *Chlamydomonas reinhardtii* (*Chlamydomonas*) contain genes for both (5), although *c*_6A_ is expressed at a low level compared to *c*_6_ or plastocyanin (6, 7). Since its discovery, *c*_6A_ has been hypothesized to have various functions, including replacement of plastocyanin (3), acting as a safety valve for excess electrons of photosynthesis (7), regulation of generation of reactive oxygen species (ROS) in PSI (8), and being involved in high light signalling (9) or thiol-based redox regulation (10, 11). However, the function of cytochrome *c*_6A_ remains elusive (7).

**Figure 1:**
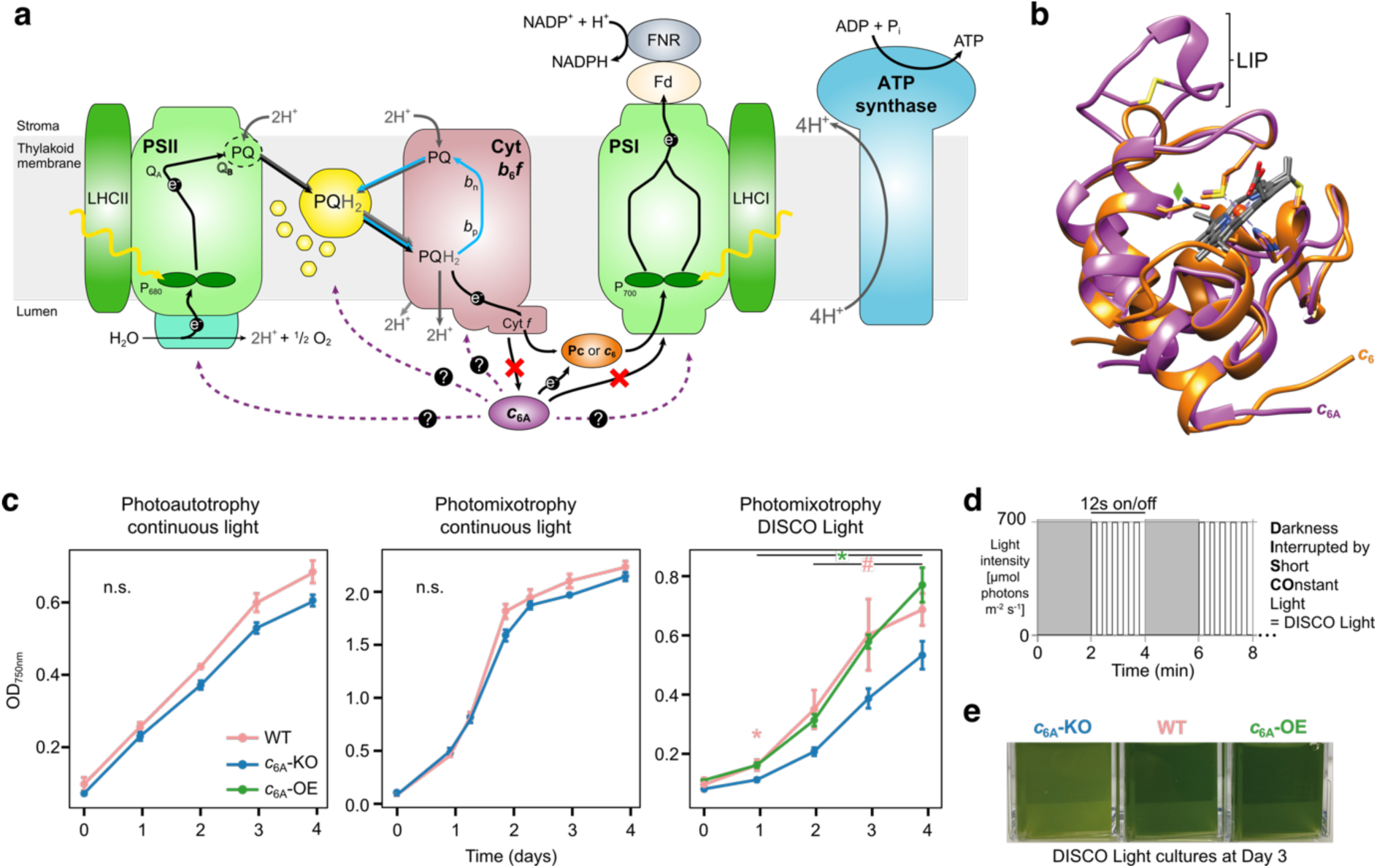
Cytochrome *c*_6A_ confers a growth advantage under a fluctuating light regime. **a**, Schematic of the photosynthetic electron transport chain with known (black) and putative interactions (purple) of *c*_6A_ marked. Proton movement (grey) and the Q-cycle (blue) are shown as arrows. **b**, Overlay of the crystal structures of cytochrome *c*_6_ from *Chlamydomonas* (orange, PDB: 1CYJ (57)) and *c*_6A_ from *A. thaliana* (purple, PDB: 2CE0 (19)). The position of the LIP is noted with the disulfide bridge formed between the two conserved cysteine residues (yellow sticks). The amino acid residues at position 52 were found to affect the heme environment contributing to the low midpoint potential of *c*_6A_ compared to *c*_6_ or plastocyanin (stick model and labelled with a green diamond) (15). **c**, Optical density (OD_750nm_) growth curve of *c*_6A_ mutant strains and WT under continuous light (30 µmol_photons_ m^-2^ s^-1^) in either photoautotrophy or -mixotrophy and DISCO Light and photomixotrophy (n = 3 biological replicates ±SD). Statistical testing was carried out with repeated measures ANOVA. Post hoc pairwise two-sided t-tests at each time point were performed if the interaction effect of time and strain on OD_750nm_ in the ANOVA was significant (p_time:strain_ < 0.05) comparing WT (pink) or *c*_6A_-OE (green) against *c*_6A_-KO (p < 0.05: *, p ≤ 0.1: #). **d**, DISCO Light (Darkness Interrupted by Short COnstant Light) consists of alternating 2 min darkness followed by 2 min of high light pulses (12 s high light (700 µmol_photons_ m^-2^ s^-1^) & 12 s darkness). **e**, Visual differences of the *c*_6A_ mutant and WT cultures of at day 3 of growth in DISCO Light.

While *c*_6_ and *c*_6A_ are homologues, there are structural differences between the two (Fig. 1b). *c*_6A_ contains a ‘loop insertion peptide’ (LIP), an insertion with a conserved disulfide bridge in a surface-located and well-defined loop. The LIP is absent from *c*_6_ (10). Assessing the stability of various mutant proteins showed that the LIP is structurally important (12, 13). While a catalytic role for the conserved disulfide bridge within the LIP is uncertain, it is predicted to contribute to the protein’s structural stability (13, 14). In addition to the LIP, there are differences in the heme environment of the two proteins, and these contribute to *c*_6A_ (+71 mV (15)) having a 254 mV lower midpoint potential than plastocyanin or *c*_6_ (+325 mV (15)).

Phylogenetic analyses revealed that *c*_6A_ represents a monophyletic group distinct from *c*_6_ and likely originates from cyanobacterial cytochrome *c*_6B_, another homologue of *c*_6_ that has a similar midpoint potential to *c*_6A_ but lacks the LIP (16). The sequence of *c*_6A_ is highly conserved across the green lineage, as with plastocyanin, suggesting an important role for *c*_6A_ within photosynthetic eukaryotes (16). However, the large difference in midpoint potentials renders it highly unlikely that *c*_6A_ acts as a functional replacement for plastocyanin in plants and algae, as *c*_6A_ would not be expected to accept electrons from the cytochrome *b*_6_*f* complex (Fig. 1a). Further, plastocyanin was found to be indispensable even when *c*_6A_ was overexpressed in *A. thaliana* (1). Additionally, the surfaces of *c*_6_ and plastocyanin are both strongly negatively charged, whereas *c*_6A_ has a strongly positive surface charge (17). Together with the difference in isoelectric points (plastocyanin: 5.1, *c*_6A_: 4.1 in *A. thaliana* (18)) this indicates different interactions.

Interactions involving *c*_6A_ further highlight its distinct role (Fig. 1a). It cannot effectively reduce PSI *in vitro*, making direct interaction between *c*_6A_ and PSI unlikely (17). Additionally, *c*_6A_ has been shown to reduce plastocyanin *in vitro* (19) and weak interaction (K_d_ = 2.01 mM) was detected with plastocyanin using NMR titrations (6), consistent with a transient interaction. Yeast Two-Hybrid assays also revealed interactions of *c*_6A_ with itself (20), the lumen thiol oxidase LTO1 (21), and the redox-regulated immunophilin FKBP13 (3). LTO1 has been reported to be involved in regulating state transitions via the kinase STT7 (22). Interaction with LTO1 and FKBP13 therefore suggests a luminal localization for *c*_6A_, consistent with its detection in the soluble fraction of the thylakoid lumen in *A. thaliana* (23). In *Chlamydomonas*, however, only a chloroplastic localization has been confirmed (6). In *A. thaliana*, *c*_6A_ was also found to be uniquely localized to sensory plastids in the epidermis and vascular parenchyma rather than mesophyll chloroplasts (24).

Photosynthetic organisms require flexibility in their photosynthetic apparatus because they grow under a diverse set of environmental conditions. These include varying light conditions from diurnal cycles to more rapid fluctuations caused by the sun, clouds, canopy, and in the case of *Chlamydomonas*, also soil and water movements. For *Chlamydomonas*, exposure to a 12 s on/12 s off cycle with light intensities of 600-700 µmol photons m⁻² s⁻¹ in photoautotrophic conditions resulted in growth rates reaching only about 30% of those achieved under continuous illumination (25). Additionally, a growth penalty was revealed under fluctuating light with frequencies in the range of 10-100 s on/off (26). In mutant assays using *Chlamydomonas*, various fluctuating light regimes have been employed to determine the function of proteins associated with photosynthesis. These include the important proteins Proton Gradient Regulation 5 (PGR5) and Proton Gradient Regulation Like 1 (PGRL1), both implicated in cyclic electron flow, its regulation or ATP synthase regulation (27), as well as flavodiiron proteins (FLVA/B), which facilitate the Mehler-like reaction (28). Additionally, the qE quenching effector LHCSR3 (29, 30) and plastid terminal oxidase PTOX2 (31) have been implicated in the response to fluctuating light, as mutants of all of these proteins showed a growth deficit under fluctuating light conditions.

Here, we present the first in-depth analysis of the connection between *c*_6A_ and photosynthesis. We show that this highly conserved protein is important for growth under a harsh fluctuating light regime we call DISCO Light and helps maintain the redox balance of the plastoquinone (PQ) pool through its effect on the light harvesting balance between photosystems I (PSI) and II (PSII).

## Results

### Cytochrome *c*_6A_ confers a growth advantage under a fluctuating light regime

To investigate the role of *c*_6A_ in *Chlamydomonas* we used CRISPR/Cpf1 (32) to create a knock-out of the *c*_6A_ gene (*CYC4*, Cre16.g670950) in the background strain CC-1883 (WT). The knock-out strain (“*c*_6A_-KO”) was confirmed through PCR and sequencing (Supplementary Methods & Supplementary Fig. 1a-d). We also generated a *c*_6A_ overexpression *c*_6A_-OE) strain by transforming the *c*_6A_-KO strain with a construct that drives the expression of FLAG-tagged *c*_6A_ from a *PSAD* promoter. Overexpression was confirmed by immunoblot and RNAseq analysis (Supplementary Methods & Supplementary Fig. 1d-e).

To investigate the role of cytochrome *c*_6A_ in growth, we tested how *Chlamydomonas* strains grow under various light and trophic regimes. Under continuous light (30 μmol photons m⁻² s⁻¹), both *c*_6A_-KO and WT strains reached similar maximum optical densities (OD_750nm_) of around 2 in photomixotrophic and ∼0.6 in photoautotrophic conditions, with no statistically significant differences (Fig. 1c). We defined continuous light (30 μmol photons m⁻² s⁻¹) and photomixotrophic conditions as standard conditions for this study.

We also tested multiple stressful light conditions for their effect on *c*_6A_-KO and WT. Only a light regime we named DISCO (Darkness Interrupted by Short COnstant) Light and photomixotrophy led to an observable phenotypic growth difference (Fig. 1c). DISCO Light was devised to expose *Chlamydomonas* to a complex fluctuating light regime covering multiple frequencies, as it consists of two alternating phases: 2 min of darkness followed by 2 min of alternating 12 s high light pulses of 700 μmol_photons_ m^-2^ s^-1^ and 12 s darkness (Fig. 1d). *c*_6A_-KO grew more slowly and reached lower cell densities than WT under photomixotrophic DISCO conditions (Fig. 1c & e): after one day *c*_6A_-KO cultures showed significantly reduced growth (*P* = 0.029), and by day 3 they were approximately 1.5 times less dense than WT cultures (*P* = 0.10). The growth penalty in *c*_6A_-KO was recovered by overexpressing *c*_6A_, with *c*_6A_-OE showing significantly higher cell densities than *c*_6A_-KO from day 1 (*P* = 0.024, Fig. 1c & e). In the DISCO light regime without acetate in the medium (photoautotrophy), no strain could grow (Supplementary Fig. 2e).

In photomixotrophy and darkness or continuous high light at the same light intensity as in the DISCO regime (700 μmol_photons_ m^-2^ s^-1^) WT and *c*_6A_-KO grew at similar rates (Supplementary Fig. 2a-b). Furthermore, fluctuating light regimes corresponding to elements of the DISCO protocol (“2 min high light & 2 min darkness”, and “12 s high light & 12 s darkness”), showed only small differences between WT and *c*_6A_-KO that were not part of a consistent trend under either photomixotrophic or photoautotrophic conditions (Supplementary Fig. 2c-d).

Measurements of total chlorophyll concentration and acetate consumption confirmed the phenotypic difference between the strains observed from OD_750nm_ measurements in both standard and DISCO Light conditions (Supplementary Fig. 2f-h). The correlation between total chlorophyll concentration and optical density was similar between WT and *c*_6A_-KO. This indicated no difference in chlorophyll concentration per cell between the strains under standard conditions or DISCO Light (Supplementary Fig. 2g). Additionally, measurements of acetate concentration in the growth medium indicated that *c*_6A_-KO consumed less acetate, likely because of lower cell densities when exposed to DISCO Light, with differences apparent from day 3 (Supplementary Fig. 2h). These results indicate that *c*_6A_ is important for growth of *Chlamydomonas* under the high intensity and rapidly fluctuating DISCO Light regime and acetate availability in photomixotrophic conditions.

### *c*_6A_ is likely localized in the thylakoid lumen and does not have strongly bound interaction partners

*c*_6A_ was detected in the soluble fraction of the thylakoid lumen in *A. thaliana* (23). In *Chlamydomonas*, however, only chloroplastic localization has been confirmed (6). To determine the subcellular localization of *c*_6A_ in *Chlamydomonas* further, we performed immunofluorescence microscopy on the *c*_6A_-FLAG overexpressing strain (*c*_6A_-OE) and its background strain (*c*_6A_-KO) grown under standard conditions. In the negative control (*c*_6A_-KO), the AlexaFluor488 signal corresponding to FLAG was weak and non-specific, i.e. appearing throughout the cell including in the flagella (Supplementary Fig. 3a). In contrast, the *c*_6A_-FLAG overexpressing sample (*c*_6A_-OE) showed a more intense (2 to 3-fold pixel intensity) and more localized fluorescence signal, confirming chloroplast-specific localization of *c*_6A_ (Supplementary Fig. 3b). Two cells covering the variation of AlexaFluor488 signal intensity were chosen as representatives to capture cell to cell differences that could arise from technical variation from fixing and staining or *c*_6A_-FLAG expression level differences between cells. Nevertheless, the absence of AlexaFluor488 signal from cell compartments other than the chloroplast in *c*_6A_-OE compared to *c*_6A_-KO supports the FLAG-specificity of the signal. The detection of anti-FLAG signal in structures resembling pyrenoid-traversing thylakoids supports a thylakoid luminal localization of the soluble *c*_6A_ (Supplementary Fig. 3c-d). While it cannot be excluded that *c*_6A_ is associated with the thylakoid membrane on the stromal side, the structure of *c*_6A_ (Fig. 1b) gives no indication of membrane-association.

To detect possible covalent or strongly bound interaction partners, we performed co-immunoprecipitation (co-IP) analyses of the *c*_6A_-FLAG overexpressing strain (*c*_6A_-OE) and the untransformed background strain (*c*_6A_-KO) using anti-FLAG-agarose beads. Immunodetection of FLAG in the precipitate revealed bands at ∼15 kDa in *c*_6A_-OE but not in *c*_6A_-KO, corresponding to mature *c*_6A_-FLAG (red arrows in Supplementary Fig. 4), with additional FLAG signals that were not *c*_6A_-FLAG specific in the 25-35 kDa range (black arrows in Supplementary Fig. 4). Under both standard (Supplementary Fig. 4c) and DISCO Light (Supplementary Fig. 4f) conditions, the immunoprecipitate did not reveal any bands unique to *c*_6A_-OE except the *c*_6A_-FLAG band, indicating that no specific interaction partners of *c*_6A_ were detected.

The combined immunofluorescence microscopy and co-IP results indicate that *c*_6A_ is likely to be localized in the thylakoid lumen, as for *A. thaliana* (23) and are consistent with any hypothesized function relying primarily on weak or transient interactions.

### *c*_6A_ is important to maintain the light harvesting balance between photosystems I and II in photomixotrophy

Given the light-dependent growth phenotype observed in DISCO conditions, the likely localization of *c*_6A_ in the thylakoid lumen, and *c*_6A_’s high sequence similarity to *c*_6_ which is involved in the PETC, we investigated the photosystems and light harvesting complexes to explore further a potential role of *c*_6A_ in photosynthesis.

We recorded low temperature (77K) chlorophyll fluorescence emission spectra. When grown in continuous light (30 µmol_photons_ m^-2^ s^-1^) under photoautotrophic conditions WT and *c*_6A_-KO exhibited very similar fluorescence spectra (Fig. 2a, PSI/PSII fluorescence ratios 1.25 and 1.3, respectively, *P* = 0.374), consistent with similar growth (Fig. 1c). However, when grown in photomixotrophic conditions and continuous light (30 µmol_photons_ m^-2^ s^-1^, standard conditions), and despite the two strains showing similar growth (Fig. 1b), WT maintained a PSI/PSII fluorescence ratio of about 1.2, while *c*_6A_-KO showed a significantly lower ratio (0.96, *P* = 0.008), which was accompanied by a more prominent CP47-associated peak at 695 nm (33) (Fig. 2a). The PSI/PSII fluorescence ratio was restored to nearly WT levels in the *c*_6A_-OE strain (Fig. 2a). When samples were grown in DISCO Light and photomixotrophy, the differences seen in standard conditions were amplified. By day 4, the PSI/PSII fluorescence ratio in *c*_6A_-KO dropped drastically to 0.66 in comparison to WT and *c*_6A_-OE (1.05 and 1.33, respectively & *P* = 0.027 and 0.0005, respectively). Interestingly, WT showed a shoulder peak at ∼681 nm (Fig. 2a), which may reflect partial detachment of light harvesting complex II (LHCII) from PSII (34). The lower level of normalized PSI fluorescence in *c*_6A_-KO in all photomixotrophic samples did not originate from lower PSI protein levels, as evidenced by PSI subunit immunoblot analyses (Supplementary Fig. 5).

**Figure 2:**
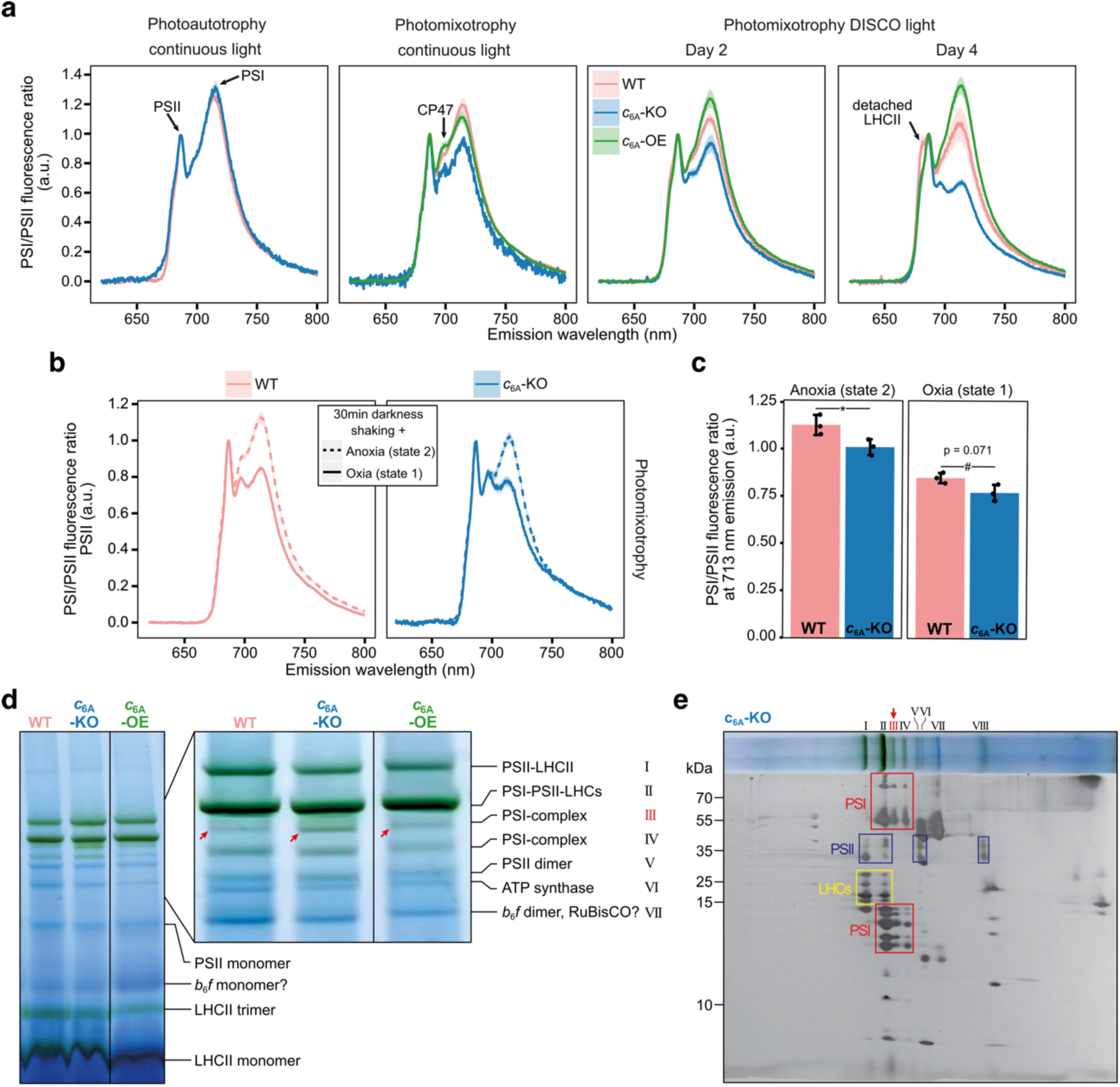
*c*6A affects the light harvesting balance between photosystems I and II in photomixotrophy. **a**, Low temperature (77 K) chlorophyll fluorescence spectra (excitation: 440 nm) of *c*_6A_ mutant strains and WT from cells grown under continuous light (30 µmol_photons_ m^-2^ s^-1^) in photoautotrophy -or mixotrophy and cells grown photomixotrophically under DISCO Light for 2 or 4 days. The prominent PSI and PSII fluorescence peaks of note are indicated by black arrows. **b**, Low temperature (77 K) chlorophyll fluorescence spectra of *c*_6A_-KO and WT exposed to state 1 or 2 inducing conditions (n = 3 biological replicates ±SE). **c**, Quantification of the PSI to PSII fluorescence ratio at 713 nm from **b**. Statistical testing was carried out with a two-sided t-test per state (p < 0.05: *, p < 0.1: #). **d**, BN-PAGE analysis of *c*_6A_ mutant strains and WT thylakoid membrane complexes isolated from cells grown in photomixotrophy and continuous light (30 µmol_photons_ m^-2^ s^-1^). The inset shows the area of interest with band III from a separate longer BN-PAGE run with the same samples. The full gels with three biological replicates per strain can be found in Supplementary Figure 6. Thylakoid membrane complexes were assigned to bands according to the literature (53–55). **e,** 2D-PAGE analysis of a BN-PAGE lane from *c*_6A_ -KO with the boxes indicating assignment of spots to protein complexes based on comparison with calculated R*_f_* values of known thylakoid membrane complex components (Supplementary Fig. 7c).

To investigate if this *c*_6A_-dependent light harvesting phenotype was due to alterations in state transitions in photomixotrophy, for example due to *c*_6A_-KO being locked in state 1, we induced either state 1 (darkness and oxia) or state 2 (darkness and anoxia) in samples grown under standard conditions (Fig. 2b). Both strains showed, as expected, a lower PSI/PSII fluorescence ratio in state 1 and a higher PSI/PSII fluorescence ratio in state 2, reflecting the increased sizes of the antenna cross section of PSII and PSI, respectively. However, *c*_6A_-KO consistently exhibited a lower PSI/PSII fluorescence ratio than WT (Fig. 2c). Whereas WT exhibited PSI fluorescence of 1.13 (normalized to PSII fluorescence) in state 2 conditions, *c*_6A_-KO showed a significantly lower value of 1.01 (*P* = 0.046). Similarly, under state 1 inducing conditions, normalized PSI fluorescence in WT was 0.85, compared to 0.77 in *c*_6A_-KO (*P* = 0.071). While the CP47-associated peak at 695 nm (33) showed different intensity between the two states in WT, in *c*_6A_-KO the same peak intensity was recorded in both states.

As the light harvesting balance is affected by photosystem complex abundance and composition, we next analyzed the thylakoid membrane complexes from cells grown in standard conditions using BN-PAGE. Gels showed a distinct green band containing a PSI complex without LHCs (band III marked red, Fig. 2d&e) that was consistently more prominent in *c*_6A_-KO compared to WT and *c*_6A_-OE. The other bands showed a similar pattern and thus presumably similar photosystem complex levels across strains (Fig. 2d). 2D-PAGE analyses and gel simulations to calculate the predicted mobility in the gel (R*_f_*) revealed similar subunit compositions of the complexes between strains (Fig. 2e & Supplementary Fig. 7a-b).

Together, these results show that *c*_6A_ is important for maintaining the light harvesting balance between the photosystems in photomixotrophic conditions, especially in combination with DISCO light, and indicate that an altered distribution of LHCs could underly this observation.

### Absence of *c*_6A_ leads to a more reduced PQ pool

Given the tight link between light harvesting balance and the redox state of the PQ pool, we quantified the redox state of the PQ pool using chlorophyll fluorescence measurements.

Firstly, the chlorophyll fluorescence induction (OJIP) traces of cells grown in standard conditions revealed a 10% increase in the F_J_ parameter in *c*_6A_-KO compared to WT and *c*_6A_-OE (*P* = 0.012 and 0.008, respectively) (Fig. 3a,b). Since F_J_ reflects the extent of Q_A_ reduction and is positively correlated with PQ pool reduction (35), this increase indicates that the absence of *c*_6A_ led to a more reduced PQ pool.

**Figure 3:**
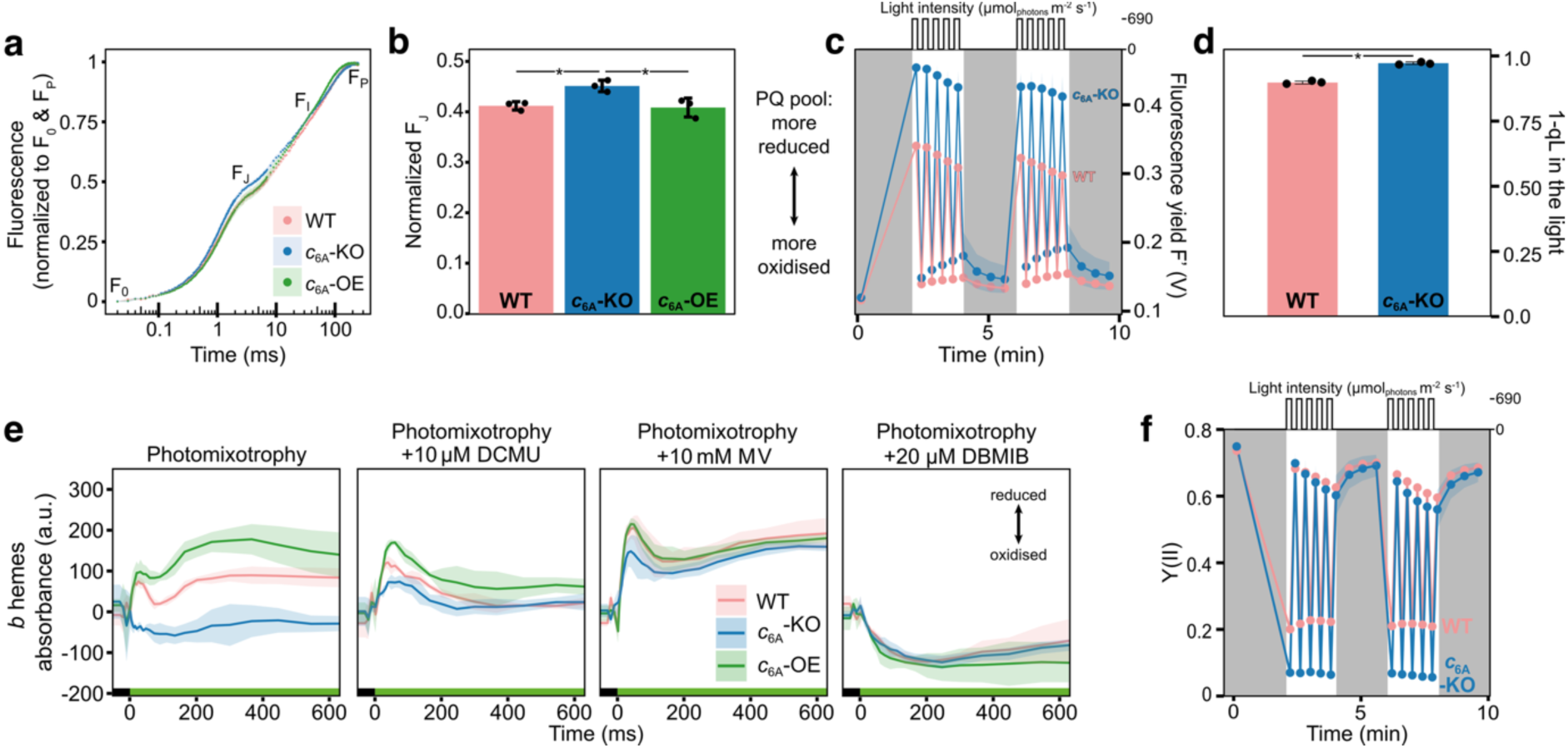
*c*_6A_ affects the redox state of the PQ pool, the redox kinetics of *b* hemes in cytochrome *b*_6_*f* and the quantum yield of PSII. **a**, Chlorophyll fluorescence induction measurements (OJIP curves) of *c*_6A_ mutant strains and WT grown under photomixotrophic continuous light (30 µmol_photons_ m^-2^ s^-1^) conditions and normalized to F_0_ and F_P_ (n = 3 biological replicates ±SD). **b**, Comparison of the normalized fluorescence at inflection point F_J_ (2.4 ms) from **a**. Asterisks indicate a significant difference (p < 0.05) as determined by one-way ANOVA and post-hoc pairwise two-sided t-test (n = 3 biological replicates ±SD). **c**, Raw steady-state fluorescence yield F’ in Volts (V) of *c*_6A_-KO and WT grown under standard conditions and exposed to a DISCO Light regime (on top of box) in a DUAL-PAM-100 spectrometer (n = 3 biological replicates ±SD). **d**, 1-qL values in the light phases of *c*_6A_-KO and WT grown under standard conditions and exposed to the same light protocol as in **c** (n = 3 biological replicates ±SD, asterisk: significant difference (p < 0.001) as determined by two-sided t-test). **e**, Redox changes of *b* hemes in *c*_6A_ mutant strains and WT grown under standard conditions and exposed to DCMU (competitive inhibitor of the Q_B_ site in PSII), methyl viologen (MV, electron acceptor of PSI) or dibromothymoquinone (DBMIB, *b*_6_*f* Q_P_ site blocker). Samples were exposed to green light (500 µmol_photons_ m^-2^ s^-1^, green bar) in a JTS-10 spectrometer (n = 3 biological replicates ±SD). **f**, Photosystem II quantum yield, Y(II), of *c*_6A_-KO and WT grown under standard conditions and exposed to the same light protocol as in **c** (n = 3 biological replicates ±SD).

To investigate further the extent of *c*_6A_-dependent PQ pool overreduction, we exposed cells grown in standard conditions to DISCO light and recorded chlorophyll fluorescence. The value of the steady-state fluorescence yield F’ under actinic light is influenced by the redox state of the PQ pool, as the PQ pool is in a redox equilibrium with Q_A_ in PSII (36). During the high light periods of DISCO Light, F’ was on average 36% higher in *c*_6A_-KO than in WT (*P* = 0.0006), indicating a more reduced PQ pool in the absence of *c*_6A_ in DISCO Light as well (Fig. 3c). Also during the high light periods of the DISCO Light regime, 1-qL (a measure of Q_A_ reduction) was high in WT (0.9) indicating that Q_A_ and by implication the PQ pool was highly reduced (37). In *c*_6A_-KO 1-qL was significantly higher (0.975) again compared to WT (*P* = 6.8 × 10^-5^) (Fig. 3d), indicating a more reduced PQ pool, consistent with the F’ finding.

These results suggest that the absence of *c*_6A_ leads to a more reduced PQ pool in *Chlamydomonas* under photomixotrophic conditions, exacerbated by DISCO Light exposure. Consequently, the more reduced Q_A_ pool could lead to increased damage to PSII, potentially contributing to the observed growth penalty in DISCO Light.

### Redox kinetics of *b* hemes are affected and PSII quantum yield lowered in absence of *c*_6A_

Given the effects of the absence of *c*_6A_ on light harvesting and PQ pool redox balance, we also assessed the function of PETC complexes. We first measured the redox kinetics of cytochrome *b*_6_*f* using a Joliot-type spectrometer (JTS).

In photoautotrophic samples grown under continuous light (30 µmol_photons_ m^-2^ s^-1^), all strains exhibited a *b* hemes reduction peak within 100 ms of light exposure, but in *c*_6A_-KO the peak was only half the intensity of WT (Supplementary Fig. 8). In samples grown in standard photomixotrophic conditions, the *b* hemes in WT and *c*_6A_-OE were reduced within 100 ms of light exposure followed by transient oxidation and re-reduction. In contrast, the *b* hemes in *c*_6A_-KO were *oxidized* within 100 ms of illumination (Fig. 3e) and showed nearly no re-reduction in the following 500 ms. Interestingly, no differences in the cytochrome *f* redox kinetics were observed in either photoautotrophy or photomixotrophy (Supplementary Fig. 8), indicating no effect of the absence of *c*_6A_ on the function of the high potential chain.

A more reduced PQ pool leads to lower reduction and even apparent oxidation of the *b* hemes upon light exposure (38–40). The latter is likely due to *b* hemes being pre-reduced in conditions with a highly reduced PQ pool. Thus, the effect of the absence of *c*_6A_ on the redox kinetics of *b* hemes are consistent with the observed change in the redox state of the PQ pool. Artificial modulation of the PQ pool redox state should therefore lead to smaller differences in *b* hemes redox kinetics between the strains. In the presence of dichlorophenyl-dimethylurea (DCMU), a PSII inhibitor that leads to PQ pool oxidation, samples grown in standard conditions showed that the *b* hemes in *c*_6A_-KO were indeed reduced upon illumination but to a lower magnitude than in WT/ *c*_6A_-OE (Fig. 3e). Similarly, when we added methyl viologen (MV), an artificial electron acceptor from PSI that also leads to PQ pool oxidation, the *b* hemes reduction was partially restored in in *c*_6A_-KO (Fig. 3e). Furthermore, when we added dibromothymoquinone (DBMIB), which blocks the Q_P_ site of cytochrome *b*_6_*f* and leads to PQ pool reduction, the *b* hemes only showed oxidation upon illumination without differences between the strains (Fig. 3e).

The observed differences in *b* hemes redox kinetics can therefore be explained by the more reduced PQ pool in *c*_6A_-KO. Further, these results indicate that in the absence of *c*_6A_ the cytochrome *b*_6_*f* complex including the Q-cycle is in principle functional if the PQ pool is oxidized. Nevertheless, small differences between *c*_6A_-KO and WT/ *c*_6A_-OE remained when the PQ pool was artificially oxidized, warranting further investigation of the Q-cycle.

To investigate the possibility of a disturbed Q-cycle underlying the *b* hemes phenotype, we measured the proton motive force (*pmf*) and its components of *Chlamydomonas* cells grown in standard conditions using electrochromic shift (ECS) measurements. No substantial difference between presence and absence of *c*_6A_ was detected in *pmf*, proton flux (vH^+^) and thylakoid conductivity (gH^+^) when samples were exposed to either continuous red light (53 µmol_photons_ m^-2^s^-1^) or DISCO Light conditions (Supplementary Fig. 9a-f). Further, no differences between the strains were found in the response of the ECS signal to a single turnover flash and the partitioning of *pmf* into ΔpH and ΔΨ (Supplementary Fig. 9g&h). Together, this is consistent with *c*_6A_ not affecting Q-cycle function.

We also analyzed the efficiency of photosystems I and II of *Chlamydomonas* cells grown in standard conditions when exposed to continuous or DISCO Light. In continuous light, we did not find any significant differences between WT and *c*_6A_-KO in the quantum yield of PSII (Supplementary Fig. 10a) and PSI (Supplementary Fig. 10b) over a wide range of intensities. There was, however, an elevated photosystem II quantum yield Y(II) for *c*_6A_-OE in comparison to *c*_6A_-KO at higher light intensities (above 210 μmol_photons_ s^-1^ m^-2^) resulting in an increased relative electron transport rate (rETRII). Conversely, during the high light periods of DISCO Light, Y(II) was less than half in *c*_6A_-KO compared to WT (Fig. 3f). In contrast to PSII, PSI quantum yield Y(I) was not majorly affected by absence of *c*_6A_ and DISCO Light and showed a decreasing trend in the light phases from 0.2 to about 0.125 over the course of 40 min in both WT and *c*_6A_-KO (Supplementary Fig. 11). For both strains, DISCO Light led to an increase in acceptor side limitation Y(NA), indicating that in DISCO Light downstream electron acceptors are not readily available (Supplementary Fig. 11). Consistent with the more reduced PQ pool in absence of *c*_6A_, *c*_6A_-KO exhibited a lower donor-side limitation Y(ND) than WT during DISCO Light treatment (Supplementary Fig. 11).

The DISCO Light condition therefore seems to lead to inhibition of photosynthesis at PSII in *c*_6A_-KO which is consistent with the highly reduced PQ pool in absence of *c*_6A_.

### *c*_6A_ partly alleviates the stressful effect of DISCO Light on PSII

To determine if the more reduced PQ pool in the absence of *c*_6A_ causes stress for *Chlamydomonas* and could explain the observed growth phenotype, we measured PSII energy partitioning of cells grown in standard conditions and then exposed to DISCO Light using a DUAL-PAM.

Increased stress levels in the absence of *c*_6A_ were evident when examining the quantum yield of non-regulated dissipation of excitation energy, Y(NO), which describes the fraction of energy that is neither used for photochemistry nor dissipated as heat in a regulated manner via NPQ. At the start of DISCO Light exposure, over 90% of light energy was dissipated in a non-regulated fashion in *c*_6A_-KO, while WT had values below 80% (Fig. 4a). Over 40 minutes of DISCO Light, Y(NO) levels in *c*_6A_-KO relaxed, nearing WT levels of about 70%. High Y(NO) levels indicate PSII photoinhibition that is imminent or has already occurred due to over-reduction (41). When probing for PsbA, C-terminal fragments forming two bands at around 15 kDa were visible (Fig. 4b). An increased ratio of PsbA (D1) degradation products to full length PsbA (D1) protein, which is indicative of PSII photoinhibition (42), was observed in *c*_6A_-KO on Day 4 of DISCO Light exposure (Fig. 4b & Supplementary Fig. 12). In *c*_6A_-KO, 20% of the observed PsbA (D1) protein signal originated from the C-terminal fragment bands, while in WT and *c*_6A_-OE the C-terminal fragments only contributed 8% and 7% to the total PsbA (D1) signal, respectively (Supplementary Fig. 12). The yield of regulated non-photochemical quenching Y(NPQ) was negligible in dark-adapted WT and *c*_6A_-KO samples (Fig. 4b). However, over the first 40 min of DISCO Light exposure, *c*_6A_-KO exhibited approximately twice the Y(NPQ) levels as WT during all phases of the DISCO Light protocol. This sustained Y(NPQ) response, even in the dark, indicates increased stress levels in the absence of *c*_6A_, leading to an amplified engagement of NPQ mechanisms to allow energy dissipation in a regulated manner. Interestingly, increased Y(NPQ) in the absence of *c*_6A_ was also observed under a range of continuous light intensities, which was partially restored in the *c*_6A_-OE strain (Supplementary Fig. 10a). Elevated NPQ levels in the absence of *c*_6A_ were further supported by an approximately 1.5-fold protein level of the NPQ effector LHCSR3 (43) in *c*_6A_-KO compared to WT after two days of growth in DISCO Light (Fig. 4b & Supplementary Fig. 12).

**Figure 4:**
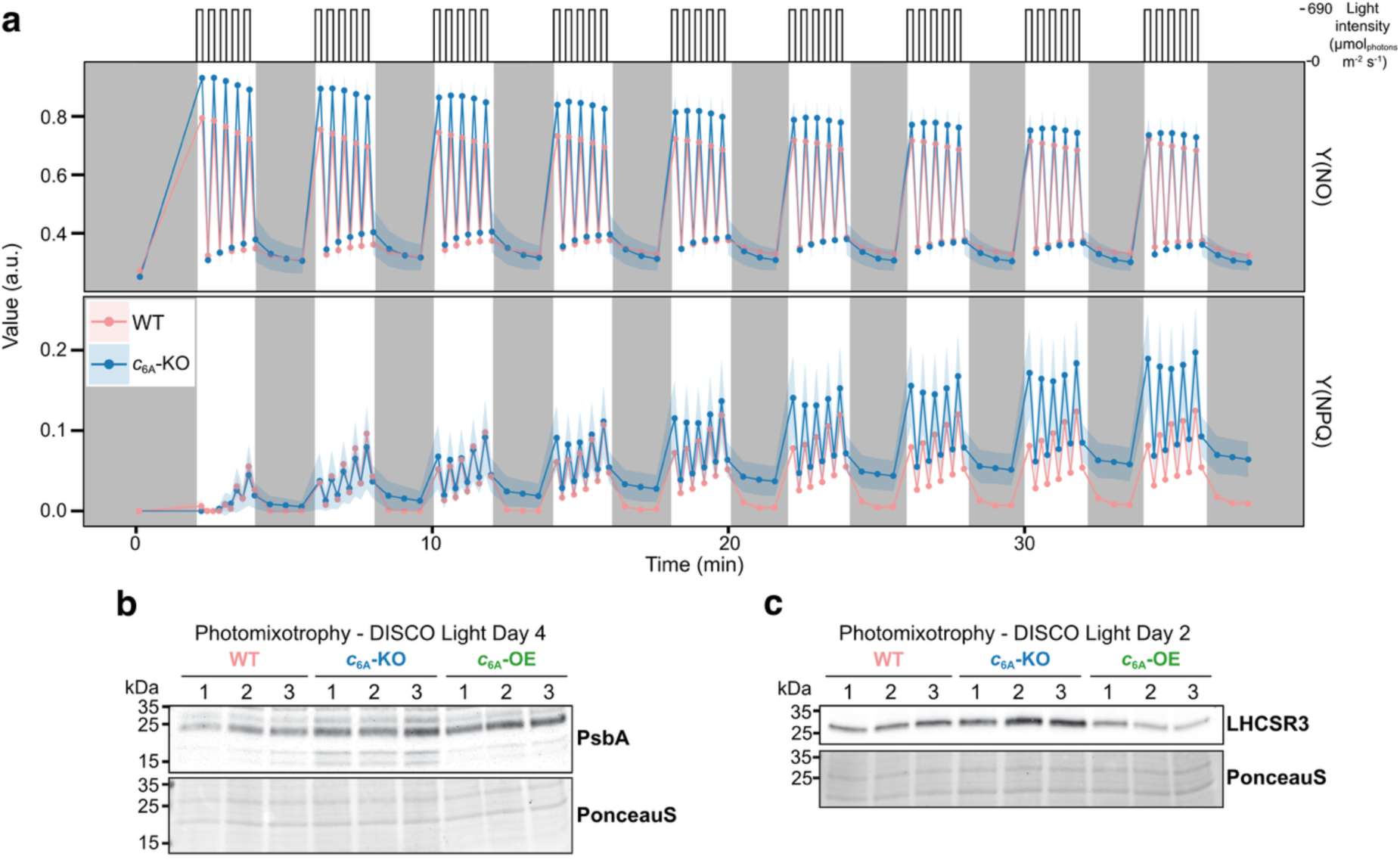
*c*_6A_ partly alleviates the stressful effect of DISCO Light on *Chlamydomonas*. **a**, Regulated non-photochemical quenching Y(NPQ), and non-regulated energy dissipation Y(NO) of *c*_6A_-KO and WT grown under standard conditions and exposed to a DISCO Light regime in a DUAL-PAM-100 spectrometer (n = 3 biological replicates ±SD). **b**, Immunoblot analyses probing for PsbA in whole cell protein samples from *c*_6A_ mutant strains and WT grown in DISCO Light for four days (n = 3 biological replicates, PonceauS stain as total protein loading control). **c**, Immunoblot analyses probing for LHCSR3 in whole cell protein samples from *c*_6A_ mutant strains and WT grown in DISCO Light for two days (n = 3 biological replicates, PonceauS stain as total protein loading control). Relative quantification of the share of signal from C-terminal fragments for PsbA or the signal intensities normalized to total protein for LHCSR3 and can be found in Supplementary Figure 12.

Together, these data indicate that PSII experiences more photoinhibition in *c*_6A_-KO than WT and *c*_6A_-OE under DISCO Light consistent with the observed reduced Y(II) in *c*_6A_-KO under DISCO Light (Fig. 3f). Increased photoinhibition is likely contributing to the observed growth penalty (Fig. 1c).

## Discussion

Here we present an in-depth characterization of the physiological role of the highly conserved cytochrome *c*_6A_ protein in relation to photosynthesis and show that it is important for optimal growth in fluctuating light due to its effect on the light harvesting and PQ pool redox state balances of the model alga *Chlamydomonas* (summary see Supplementary Table 1).

Analysis of the redox state of the PQ pool using complementary chlorophyll fluorescence approaches indicated that the PQ pool was more reduced in *c*_6A_-KO, and this was particularly marked in the high light periods of DISCO Light. JTS measurements of the redox state of the *b* hemes of the cytochrome *b*_6_*f* complex in response to light exposure showed disturbance of the redox kinetics in the *c*_6A_-KO that were consistent with a more reduced PQ pool in *c*_6A_-KO. Addition of DCMU or MV, which would be expected to lead to PQ oxidation, or DBMIB, which leads to PQ reduction, revealed effects consistent with an increased pre-reduction of the *b* hemes in *c*_6A_-KO through a more reduced PQ pool. *c*_6A_-KO exhibited no change in the proton motive force, measured by electrochromic shift, rendering unlikely a direct effect of *c*_6A_ on the Q cycle. Under the high light periods of DISCO Light, *c*_6A_-KO showed a dramatic decrease in PSII quantum yield compared to WT, which would also be consistent with a more reduced PQ pool. The *c*_6A_-KO did not exhibit any significant difference in PSI quantum yield under DISCO light, although it showed a decrease in donor-side limitation of PSI, again consistent with a more reduced PQ pool. *c*_6A_-KO also showed some indication of increased acceptor-side limitation.

*c*_6A_-KO exhibited an increase in the non-regulated dissipation of excitation energy [Y(NO)] and regulated non-photochemical quenching [Y(NPQ)] with the onset of DISCO Light, as well as a relative increase in the amount of the NPQ effector LHCSR3 protein and degradation products of PsbA after two and four days of growth in DISCO Light, respectively. These results indicate an increased tendency to photoinhibition in *c*_6A_-KO, which might be accounted for by the increased PQ reduction. A general impairment of PSII function or repair in the absence of any other effects from loss of *c*_6A_ might be expected to lead to a decreased reduction in PQ. This suggests that the PSII impairment is a consequence of the increased PQ reduction resulting from loss of *c*_6A_, rather than the primary result of *c*_6A_ loss.

What might cause the increase in reduction of the PQ pool in the absence of *c*_6A_? The 77K chlorophyll fluorescence data indicated a significantly lower PSI/PSII fluorescence ratio in *c*_6A_-KO compared to WT under photomixotrophic conditions, and this difference was amplified under DISCO Light conditions. In BN-PAGE, a band corresponding to a LHC-less PSI was more prominent in *c*_6A_-KO than WT and *c*_6A_-KO showed a lower PSI/PSII fluorescence value than the wild type under both state 1 and state 2 conditions. These data are consistent with the photosystems being preferentially in the equivalent of state 1 in *c*_6A_-KO compared to WT. However, there was no evidence for the *c*_6A_-KO being permanently ‘locked’ in either state 1 or state 2.

Overall, it therefore seems likely that *c*_6A_-KO is disturbed in the light harvesting balance between PSI and PSII, in favor of PSII, especially under DISCO Light. This leads to a more reduced PQ pool, and consequent photoinhibition, with impaired growth of *c*_6A_-KO in DISCO conditions. *c*_6A_ is likely to be located in the thylakoid lumen and has been shown to interact with the lumen thiol oxidase LTO1 (21), which is closely involved in thiol-based regulation of the STT7 (STN7 in Arabidopsis) state transition kinase, keeping STT7 oxidized (22). STT7 is active only when oxidized and, in response to a reduced PQ pool, catalyzes phosphorylation of LHCII and causing LHCII to move from PSII to PSI. *c*_6A_ has been proposed to catalyze the formation of disulfide bridges in the lumen (10, 11), transferring electrons to plastocyanin, with which it has been shown to react (19). Although we cannot exclude a more complex explanation, with the shift in PSI/PSII balance and change in state of the PQ pool being indirect consequences of loss of *c*_6A_, we propose that *c*_6A_ contributes to oxidation of STT7. *c*_6A_ is not essential for the oxidation as state transitions remain in *c*_6A_-KO. However, it may be that *c*_6A_ assists in the transfer of electrons from STT7 to LTO1 (with which it interacts (21)) or another acceptor. Thus, in the absence of *c*_6A_, STT7 would be more reduced than in WT, leading to increased light harvesting at PSII and a more reduced PQ pool compared to WT. This model could be tested in the future by making substitutions to the conserved cysteine residues in *c*_6A_. If correct, this will be a significant contribution to our understanding of the complex thiol-based regulatory network of the thylakoid lumen of photosynthetic organisms (44). Furthermore, the demonstration of the importance of *c*_6A_ for growth under fluctuating light conditions presents a significant contribution to understand how organisms cope with rapidly changing light conditions (45, 46). The connection of *c*_6A_ to state transitions also make it a significant target for optimizing crop productivity.

## Materials and Methods

### Growth conditions

*Chlamydomonas* strains were maintained on Tris-Acetate-Phosphate (TAP) (47) medium with Hutner’s trace element solution (48) supplemented with 1.5% agar at 30 µmol_photons_ m^-2^ s^-1^ and 25 °C. For longer-term storage cultures were kept on TAP slants in ambient light at room temperature. Liquid cultures were grown at 30 µmol_photons_ m^-2^ s^-1^, 25 °C and 120 rpm in AlgaeTron AG130 or AG230 incubators (Photon System Instruments) unless otherwise specified.

For experimental cultures, triplicates of 30 mL medium (Tris-phosphate (TP) medium (47) for photoautotrophic, and TAP for photomixotrophic conditions) were inoculated with *Chlamydomonas* strains to OD_750nm_ = 0.1. The cultures were then subjected to the experimental conditions (Table 1). All experimental cultures were subcultured at least twice in standard conditions before exposure to experimental conditions.

**Table 1:**
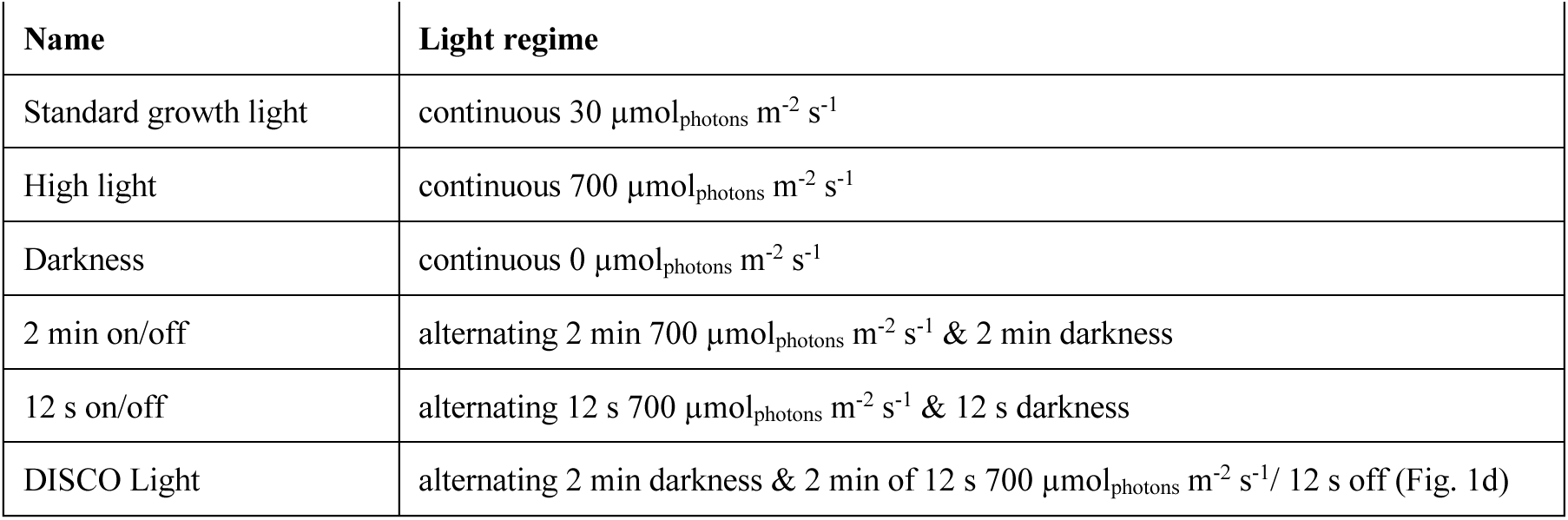
Experimental light conditions in the AlgaeTron AG130 and AG230 growth chambers.

| Name | Light regime |
| --- | --- |
| Standard growth light | continuous $30 \mu\text{mol}_{\text{photons}} \text{m}^{-2} \text{s}^{-1}$ |
| High light | continuous $700 \mu\text{mol}_{\text{photons}} \text{m}^{-2} \text{s}^{-1}$ |
| Darkness | continuous $0 \mu\text{mol}_{\text{photons}} \text{m}^{-2} \text{s}^{-1}$ |
| 2 min on/off | alternating 2 min $700 \mu\text{mol}_{\text{photons}} \text{m}^{-2} \text{s}^{-1}$ & 2 min darkness |
| 12 s on/off | alternating 12 s $700 \mu\text{mol}_{\text{photons}} \text{m}^{-2} \text{s}^{-1}$ & 12 s darkness |
| DISCO Light | alternating 2 min darkness & 2 min of 12 s $700 \mu\text{mol}_{\text{photons}} \text{m}^{-2} \text{s}^{-1}$ / 12 s off (Fig. 1d) |

### Algal strains

The cytochrome *c*_6A_ knock-out strain “*c*_6A_-KO” (CC-1883 background) was generated as previously described (32) using the gRNAs 5’-ATTTACAGTTAGTGCATCCACTTG-3’ and 5’-TCCGAGGGTAAGCAAGCACGGCAC-3’ to target the 5’UTR and Exon III for Cpf1-mediated double strand breaks, respectively. A repair template with the following sequence was provided at the same time to introduce a multiple stop codon (lowercase letters): 5’-TCATGCTCGGCCAGTTTTCTTTGATTTCAAATCCTCGCTAATTTActaattaactaaCAGCTTCCTTGCTTCATGGACCCAGTAACGGTGCATGGGTACGCG-3’.

The *c*_6A_-overexpression strain *c*_6A_-OE was created by transforming *c*_6A_-KO with the constructs pDKM-000 by electroporation as described before (49) and specified in the Supplementary Methods.

### Protein extraction and Western Blot

A known volume of *Chlamydomonas* cells at a desired growth stage was harvested by centrifugation at 1000 xg for 5 min. The pellet was subsequently resuspended in RIPA lysis buffer with freshly added PMSF (1mM final concentration) to reach a calculated OD_750nm_ of 50. The chlorophyll content was measured as above, but here 10 µL of sample were mixed with 40 µL of acetone and 1 mL of 80% acetone, resulting in a dilution factor D of 105. Protein samples were then adjusted to the same total chlorophyll concentration with RIPA buffer and frozen at -20 °C for at least 1.5 h or overnight, followed by thawing on ice to complete cell lysis. Protein concentration was determined using a Bradford assay (Merck) according to manufacturer’s instructions diluting the protein samples 5- to 10-fold.

For SDS-PAGE, all samples were diluted to 1 µg/µL protein with RIPA buffer before being mixed with 4x Lämmli buffer and incubated for 30 min (50 °C, 600 rpm). After centrifuging the samples (12000 xg, 1 min) to remove insoluble material, samples were loaded to the desired amount into wells of polyacrylamide gels (Mini-PROTEAN TGX Stain-Free, BioRad). PageRuler Plus (7 µL, Thermo Scientific) was used as size marker. Electrophoresis was carried out at 4 °C and 200 V for about 50 min. The gel was imaged in a ChemiDoc MP imaging system (BioRad) using the Stain Free setting and 5 min activation time. Some gels were stained with Pierce™ Silver Stain for Mass Spectrometry (Thermo Scientific).

Transfer of protein from polyacrylamide gels onto polyvinylidene fluoride (PVDF) membranes was carried out using the iBlot dry transfer apparatus, following the manufacturer’s instructions and using program P3 for 5 min. Membranes were then probed for successful transfer with PonceauS (Thermo Scientific) as per manufacturer’s instructions and imaged with the ChemiDoc MP imaging system (BioRad). Following the PonceauS destaining, membranes were blocked in 5% milk in Tris-buffered saline with 0.1% Tween 20 (TBS-T) for 1 h at room temperature with gentle shaking. After washing membranes once with TBS-T to remove remnants of blocking agent, membranes were incubated with specific primary antibody solutions in 5% milk TBS-T overnight at 4 °C (PsaB (AS10695, Agrisera) at 1:1000, PsbA D1 (AS05084, Agrisera) at 1:10000, LHCSR3 (AS142766, Agrisera) at 1:2000). Subsequently, membranes were washed with TBS-T (5x 5 min) and incubated for 1.5 h at room temperature with goat anti-rabbit horseradish peroxidase-linked secondary antibody in 5% milk TBS-T (1:10000, 1706515, BioRad). After washing membranes thrice in TBS-T and once in TBS (10 min each), they were incubated with chemiluminescence substrate (SuperSignal™ West Pico PLUS Chemiluminescent Substrate, Thermo Scientific) and imaged using a ChemiDoc MP imaging system (BioRad).

### Low temperature chlorophyll (77K) fluorescence

77K fluorescence emission spectra were recorded with a QE Pro spectrophotometer (Ocean Optics, Dunedin, FL, USA). Cell samples (200 µL) frozen in liquid N_2_ were excited with blue light (440 nm). Raw spectra were normalized to the photosystem II (PSII) fluorescence peak at 685 nm. *Chlamydomonas* cells were harvested and total chlorophyll concentration was adjusted to 5 µg/mL with fresh medium (TP or TAP). Cells were then frozen in liquid N_2_ for measurement. For the state transition experiment, *Chlamydomonas* cells grown in TAP under standard conditions were adjusted to 5 µg/mL total chlorophyll with fresh medium. Anoxia (state 2 inducing) samples were adjusted with medium containing glucose (final concentration 20 mM) and Glucose Oxidase (final concentration 50 U/mL) that was purged with N2 for 3 min. The final samples were again purged with N_2_ before being sealed. Oxia (state 1 inducing) samples were adjusted with unchanged medium and left unsealed. All samples were then shaken for 30 min in darkness before being frozen in liquid N_2_ for fluorescence measurement.

### Blue native PAGE and second dimension SDS-PAGE

Thylakoid membranes from *Chlamydomonas* cultures were isolated over a discontinuous Percoll gradient as described before (50). BN-PAGE was then carried out largely as previously (51). Thylakoid samples were diluted with resuspension buffer (25 mM Bis-Tris/HCl (pH 7.0), 20% (w/v) glycerol, 0.25 mg/ml Pefabloc, and 10 mM sodium fluoride) to a chlorophyll concentration of 0.53 µg/µL and to a total of 6 µg chlorophyll. An equal volume of β -dodecyl maltoside (β -DM, 2% in resuspension buffer) was added to each sample and the thylakoid membranes were solubilized by gentle pipetting for 2 min. After centrifugation (4 °C, 18000 xg, 20 min) samples were mixed with a one-tenth volume of sample loading buffer (100 mM Bis-Tris/HCl (pH 7.0), 0.5 M ACA, 30% (w/v) sucrose and 50 mg/ml Serva Blue G) and loaded into the wells of a NativePAGE 3-12% Bis-Tris protein gel (Invitrogen). Electrophoresis was carried out with specific anode buffer (50 mM Bis-Tris/HCl, pH 7.0) and cathode buffer containing Serva Blue G dye (50 mM Tricine, 15 mM Bis-Tris/HCl, pH 7.0, and 0.01% Serva Blue G) at 4 °C for 20 min at 75 V, 30 min at 100 V, 30 min at 125 V, 60 min at 150 V, 30 min at 175 V and 60 min at 200 V. After the 30 min at 125 V, the blue cathode buffer was replaced by the same cathode buffer but without Serva Blue G. The last step was prolonged to 5 h if additional separation of bands of interest was needed.

Gel electrophoretic separation of complexes in BN-PAGE lanes on a second dimension SDS-PAGE was carried out as previously described (51, 52), although 12% polyacrylamide gels were used here. Gels were stained with Pierce™ Silver Stain (Thermo Scientific).

The predicted mobility in the gel (R*_f_*) values of known thylakoid membrane complex components were calculated using their known molecular weight and a linear regression of the distance of the protein standard in relation to the running front (r^2^ = 0.98). The 2D-PAGE spots were compared with the R*_f_* values and with published BN-PAGE analyses from *Chlamydomonas* (53–55) to assign the bands and proteins.

### Chlorophyll fluorescence

Chlorophyll fluorescence induction (OJIP) of *Chlamydomonas* was measured with a FL3500 fluorometer (Photon System Instruments). Prior to measurement, cells were resuspended in TAP and total chlorophyll concentration was adjusted to 20 µg/mL. After recovery under standard growth conditions for at least 30 min, chlorophyll concentration was readjusted to 5 µg/mL. After 15 min incubation in darkness, fluorescence induction in response to red light (984 µmol_photons_ m^-2^s^-1^, 45% A-voltage) was recorded for 120 s. Raw fluorescence data was double normalized to fluorescence at time point 0 and the maximum fluorescence value to compare the kinetics between samples (37).

Chlorophyll fluorescence of *Chlamydomonas* was measured with a pulse amplitude-modulated spectrometer (Dual-PAM-100, Walz). Prior to measurements, cells were resuspended in TAP and the chlorophyll concentration was adjusted to 20 µg/mL. After recovery under standard growth conditions for at least 30 min, measurements were performed in a quartz cuvette at 25 °C with stirring and samples were initially incubated in darkness for 5 min before the blue fluorescence measuring light (Intensity 12, 28 µmol_photons_ m^-2^ s^-1^) was turned on. Saturating pulses (Intensity 5, 5000 µmol_photons_ m^-2^ s^-1^, 500 ms width) for PSII parameter determination were applied with stirring turned off. DISCO Light conditions in the DUAL-PAM-100 were achieved by alternating 2 min darkness followed by 2 min consisting of 12 s high light (690 µmol_photons_ m^-2^ s^-1^) and 12 s darkness alternations. Saturating pulses were applied in the middle of each 12 s phase or three times during the 2 min dark phases. F’ was recorded as the fluorescence yield during illumination with the actinic light treatment before application of a saturating pulse. Photosynthetic parameters were calculated as described previously (41).

### Cytochrome b_6_*f* redox kinetics

Cytochrome *f* and *b* heme redox kinetics were determined in *Chlamydomonas* cells by deconvoluting absorbance changes at 546, 554, 563, and 573 nm that were measured using a JTS-10 pump probe spectrophotometer (BioLogic) and appropriate 10 nm FWHM (full width at half maximum) interference filters (Edmund Optics). BG39 filters (Schott) were used to shield the light detectors from scattered light. Prior to measurements, cells were resuspended in fresh medium to a total chlorophyll concentration of 20 µg/mL. After recovery under standard growth conditions for at least 30 min, Ficoll 400 was added to a final concentration of 10%. For some samples, DCMU, methyl viologen, or DBMIB were added to a final concentration of 10 µM, 10 mM, or 20 µM respectively, at the same time as Ficoll. Cells were dark-adapted for 3 min before measurement with each interference filter and then illuminated with 5 s of green light (500 µmol_photons_ m^-2^ s^-1^). Flashes of white detection light were administered during 300 µs dark intervals in actinic illumination. Every recording was corrected for any detection light impact by subtracting a trace with actinic light turned off. Deconvolution was performed with the JTS-10 software by drawing a baseline between 573 and 546 nm. During analysis, the signal-to-noise ratio was improved with a Savitzky-Golay filter (56).

## Acknowledgments

We thank Dr. K. Geissler (University of Cambridge) for the provision of the pCM0 MoClo parts and Dr. J. Walter (University of Cambridge) for assistance with blue native PAGE. We are also grateful to Dr. M. Jokel (University of Turku) for discussions on cell culture. All photosynthesis experiments were performed within the PHOTOSYN infrastructure at the University of Turku. We gratefully acknowledge the support of this work by: the Gates Cambridge Trust with support from the Benn W Levy Trust, and the Professor John Glover Memorial Fund (to DK); the Novo Nordisk Foundation (PhotoCat, project no. NNF20OC0064371, to YA; and Photo-e-microbes project no. NNF22OCOO79717 to LTW); the Research Council of Finland no. 355335 to YA, no. 354876 to LN; the Biotechnology and Biological Sciences Research Council (BB/M011194/1 to AS, BB/M011194/1 to JML, BB/R506163/1 to AF, and BB/Y001370/1 to CJH), and Trinity Hall, Cambridge (to JML).

## Author Contributions

DK conceived the project, designed and performed experiments under the supervision of CJH and YA, and analyzed data. DK, LTW, LN, and YA developed the biophysical part of the project. LTW and LN supervised the biophysical measurements and assisted with data analysis and interpretation. AF created the knock-out mutant line under the supervision of AM. JML and AS assisted with data collection, analysis and interpretation. AM, YA, and CJH secured funding, assisted with data interpretation, and provided resources. DK wrote the manuscript with comments and input from all authors.

## References

1. M. Weigel, et al., Plastocyanin is indispensable for photosynthetic electron flow in Arabidopsis thaliana. J. Biol. Chem. 278, 31286–31289 (2003).

2. G. A. Peschek, C. Obinger, M. Paumann, The respiratory chain of blue-green algae (cyanobacteria). Physiol. Plant. 120, 358–369 (2004).

3. R. Gupta, Z. He, S. Luan, Functional relationship of cytochrome c6 and plastocyanin in Arabidopsis. Nature 417, 567–571 (2002).

4. J. Wastl, D. S. Bendall, C. J. Howe, Higher plants contain a modified cytochrome c6. Trends Plant Sci. 7, 244–245 (2002).

5. J. Wastl, S. Purton, D. S. Bendall, C. J. Howe, Two forms of cytochrome c6 in a single eukaryote. Trends Plant Sci. 9, 474–476 (2004).

6. I. Nimmo, Location and function of cytochrome c6A. PhD Thesis Univ. Camb. (2011).

7. C. J. Howe, R. H. Nimmo, A. C. Barbrook, D. S. Bendall, “Cytochrome c6A of Chloroplasts” in Cytochrome Complexes: Evolution Structures, Energy Transduction and Signaling, Advances in Photosynthesis and Respiration., W. A. Cramer, T. Kallas, Eds. (Springer Science+Business Media, 2016).

8. U. Hani, et al., A complex and dynamic redox network regulates oxygen reduction at photosystem I in Arabidopsis. Plant Physiol. (2024). 10.1093/plphys/kiae501.

9. B. A. Slater, The ancestry and function of cytochrome c6A. PhD Thesis Univ. Camb. (2020).

10. C. J. Howe, B. G. Schlarb-Ridley, J. Wastl, S. Purton, D. S. Bendall, The novel cytochrome c6 of chloroplasts: a case of evolutionary bricolage? J. Exp. Bot. 57, 13–22 (2006).

11. B. G. Schlarb-Ridley, R. H. Nimmo, S. Purton, C. J. Howe, D. S. Bendall, Cytochrome c 6A is a funnel for thiol oxidation in the thylakoid lumen. FEBS Lett. 580, 2166–2169 (2006).

12. H. Chida, et al., Crystal structure of oxidized cytochrome c 6A from Arabidopsis thaliana. FEBS Lett. 580, 3763–3768 (2006).

13. J. A. R. Worrall, B. F. Luisi, B. G. Schlarb-Ridley, D. S. Bendall, C. J. Howe, Cytochrome c 6A: discovery, structure and properties responsible for its low haem redox potential. Biochem. Soc. Trans. 36, 1175–1179 (2008).

14. J. M. Mason, D. S. Bendall, C. J. Howe, J. A. R. Worrall, The role of a disulfide bridge in the stability and folding kinetics of Arabidopsis thaliana cytochrome c6A. Biochim. Biophys. Acta BBA - Proteins Proteomics 1824, 311–318 (2012).

15. J. A. R. Worrall, et al., Modulation of Heme Redox Potential in the Cytochrome c 6 Family. J. Am. Chem. Soc. 129, 9468–9475 (2007).

16. B. Slater, D. Kosmützky, R. E. R. Nisbet, C. J. Howe, The Evolution of the Cytochrome c6 Family of Photosynthetic Electron Transfer Proteins. Genome Biol. Evol. 13, 1–12 (2021).

17. F. P. Molina-Heredia, et al., A new function for an old cytochrome? Nature 424, 33–34 (2003).

18. J. Wastl, et al., Redox properties of Arabidopsis cytochrome c6 are independent of the loop extension specific to higher plants. Biochim. Biophys. Acta BBA - Bioenerg. 1657, 115–120 (2004).

19. M. J. Marcaida, et al., Structure of Cytochrome c6A, a Novel Dithio-cytochrome of Arabidopsis thaliana, and its Reactivity with Plastocyanin: Implications for Function. J. Mol. Biol. 360, 968–977 (2006).

20. P. Pesaresi, et al., Mutants, overexpressors, and interactors of arabidopsis plastocyanin isoforms: Revised roles of plastocyanin in photosynthetic electron flow and thylakoid redox state. Mol. Plant 2, 236–248 (2009).

21. Y. Lu, et al., Identification of Potential Targets for Thylakoid Oxidoreductase AtVKOR/LTO1 in Chloroplasts. Protein Pept. Lett. 22, 219–225 (2015).

22. J. Wu, et al., Functional redox links between lumen thiol oxidoreductase1 and serine/threonine-protein kinase STN7. Plant Physiol. 186, 964–976 (2021).

23. P. J. Gollan, A. Trotta, A. A. Bajwa, I. Mancini, E.-M. Aro, Characterization of the Free and Membrane-Associated Fractions of the Thylakoid Lumen Proteome in Arabidopsis thaliana. Int. J. Mol. Sci. 22, 8126 (2021).

24. J. Beltrán, et al., Specialized Plastids Trigger Tissue-Specific Signaling for Systemic Stress Response in Plants. Plant Physiol. 178, 672–683 (2018).

25. M. Janssen, et al., Efficiency of light utilization of Chlamydomonas reinhardtii under medium-duration light/dark cycles. J. Biotechnol. 78, 123–137 (2000).

26. P. J. Graham, B. Nguyen, T. Burdyny, D. Sinton, A penalty on photosynthetic growth in fluctuating light. Sci. Rep. 7, 1–11 (2017).

27. L. Nikkanen, et al., PGR5 is needed for redox-dependent regulation of ATP synthase both in chloroplasts and in cyanobacteria. [Preprint] (2025). Available at: https://www.biorxiv.org/content/10.1101/2024.11.03.621747v2 [Accessed 29 March 2026].

28. M. Jokel, X. Johnson, G. Peltier, E. Aro, Y. Allahverdiyeva, Hunting the main player enabling Chlamydomonas reinhardtii growth under fluctuating light. Plant J. 94, 822–835 (2018).

29. C. J. Steen, A. Burlacot, A. H. Short, K. K. Niyogi, G. R. Fleming, Interplay between LHCSR proteins and state transitions governs the NPQ response in Chlamydomonas during light fluctuations. Plant Cell Environ. 45, 2428–2445 (2022).

30. T. Roach, LHCSR3-Type NPQ Prevents Photoinhibition and Slowed Growth under Fluctuating Light in Chlamydomonas reinhardtii. Plants 9, 1604 (2020).

31. W. J. Nawrocki, et al., Chlororespiration Controls Growth Under Intermittent Light. Plant Physiol. 179, 630–639 (2019).

32. A. Ferenczi, D. E. Pyott, A. Xipnitou, A. Molnar, Efficient targeted DNA editing and replacement in Chlamydomonas reinhardtii using Cpf1 ribonucleoproteins and single-stranded DNA. Proc. Natl. Acad. Sci. 114, 13567–13572 (2017).

33. E. G. Andrizhiyevskaya, et al., Origin of the F685 and F695 fluorescence in Photosystem II. Photosynth. Res. 84, 173–180 (2005).

34. J. Kargul, J. Nield, J. Barber, Three-dimensional reconstruction of a light-harvesting complex I-photosystem I (LHCI-PSI) supercomplex from the green alga Chlamydomonas reinhardtii: Insights into light harvesting for PSI. J. Biol. Chem. 278, 16135–16141 (2003).

35. S. Z. Tóth, G. Schansker, R. J. Strasser, A non-invasive assay of the plastoquinone pool redox state based on the OJIP-transient. Photosynth. Res. 93, 193 (2007).

36. B. A. Diner, Dependence of the deactivation reactions of Photosystem II on the redox state of plastoquinone pool a varied under anaerobic conditions. Equilibria on the acceptor side of Photosystem II. Biochim. Biophys. Acta BBA - Bioenerg. 460, 247–258 (1977).

37. H. M. Kalaji, et al., Frequently asked questions about chlorophyll fluorescence, the sequel. Photosynth. Res. 132, 13–66 (2017).

38. P. Joliot, A. Joliot, Proton pumping and electron transfer in the cytochrome bf complex of algae. Biochim. Biophys. Acta BBA - Bioenerg. 849, 211–222 (1986).

39. F. Buchert, M. Scholz, M. Hippler, Electron transfer via cytochrome b 6 f complex displays sensitivity to antimycin A upon STT7 kinase activation. Biochem. J. 479, 111–127 (2022).

40. A. Malnoë, F.-A. Wollman, C. de Vitry, F. Rappaport, Photosynthetic growth despite a broken Q-cycle. Nat. Commun. 2, 301 (2011).

41. C. Klughammer, U. Schreiber, Complementary PS II quantum yields calculated from simple fluorescence parameters measured by PAM fluorometry and the Saturation Pulse method. PAM Appl. Notes 1, 27–35 (2008).

42. W. J. Nawrocki, et al., Molecular origins of induction and loss of photoinhibition-related energy dissipation q I. Sci. Adv. 7, 1–13 (2021).

43. M. Ballottari, et al., Identification of pH-sensing Sites in the Light Harvesting Complex Stress-related 3 Protein Essential for Triggering Non-photochemical Quenching in Chlamydomonas reinhardtii. J. Biol. Chem. 291, 7334–7346 (2016).

44. D. Hoh, J. E. Froehlich, D. M. Kramer, Redox regulation in chloroplast thylakoid lumen: The pmf changes everything, again. Plant Cell Environ. 47, 2749–2765 (2024).

45. N. A. Eckardt, et al., Lighting the way: Compelling open questions in photosynthesis research. Plant Cell 14, 1–16 (2024).

46. S. P. Long, et al., Into the Shadows and Back into Sunlight: Photosynthesis in Fluctuating Light. Annu. Rev. Plant Biol. 73, 617–648 (2022).

47. D. S. Gorman, R. P. Levine, Cytochrome f and plastocyanin: their sequence in the photosynthetic electron transport chain of Chlamydomonas reinhardi. Proc. Natl. Acad. Sci. 54, 1665–1669 (1965).

48. S. H. Hutner, L. Provasoli, A. Schnatz, C. P. Haskins, Some Approaches to the Study of the Role of Metals in the Metabolism of Microorganisms. Proc. Am. Philos. Soc. 94, 152–170 (1950).

49. P. Crozet, et al., Birth of a Photosynthetic Chassis: A MoClo Toolkit Enabling Synthetic Biology in the Microalga *Chlamydomonas reinhardtii*. ACS Synth. Biol. 7, 2074–2086 (2018).

50. C. B. Mason, T. M. Bricker, J. V. Moroney, A rapid method for chloroplast isolation from the green alga Chlamydomonas reinhardtii. Nat. Protoc. 1, 2227–2230 (2006).

51. S. Järvi, M. Suorsa, V. Paakkarinen, E. M. Aro, Optimized native gel systems for separation of thylakoid protein complexes: Novel super- and mega-complexes. Biochem. J. 439, 207–214 (2011).

52. M. Rantala, V. Paakkarinen, E. M. Aro, Analysis of thylakoid membrane protein complexes by blue native gel electrophoresis. J. Vis. Exp. 2018, 1–6 (2018).

53. E. Devadasu, et al., Long- and short-term acclimation of the photosynthetic apparatus to salinity in Chlamydomonas reinhardtii. The role of Stt7 protein kinase. Front. Plant Sci. 14, 1–17 (2023).

54. J. G. García-Cerdán, et al., A thylakoid membrane-bound and redox-active rubredoxin (RBD1) functions in de novo assembly and repair of photosystem II. Proc. Natl. Acad. Sci. U. S. A. 116, 16631–16640 (2019).

55. B. Spaniol, et al., Complexome profiling on the Chlamydomonas lpa2 mutant reveals insights into PSII biogenesis and new PSII associated proteins. J. Exp. Bot. 73, 245–262 (2022).

56. M. J. E. Savitzky, A.; Golay, Smoothing and Differentiation of Data. Anal Chem 36, 1627–1639 (1964).

57. C. A. Kerfeld, H. P. Anwar, R. Interrante, S. Merchant, T. O. Yeates, The Structure of Chloroplast Cytochrome c6 at 1.9 Å Resolution: Evidence for Functional Oligomerization. J. Mol. Biol. 250, 627–647 (1995).

