## Supplementary Information for "A conserved photosynthetic cytochrome enhances growth of *Chlamydomonas reinhardtii* in fluctuating light"

### Supplementary Methods

#### Algal strains

The knock-out in the strain "$c_{6A}$-KO" was confirmed through whole locus PCR, showing a truncated locus in the $c_{6A}$-KO strain compared to WT (Supplementary Fig. 1b). Sanger sequencing validated the deletion of the *CYC4* sequence from 5'UTR to exon III and the introduction of a sequence with multiple stop codons. Whole genome sequencing with high read depth in both strains showed the $c_{6A}$ locus was disrupted in the targeted region in $c_{6A}$-KO but not in WT (Supplementary Fig. 1c). Transcriptomic analysis showed no $c_{6A}$ gene transcripts in the $c_{6A}$-KO strain, indicating effective disruption of transcription (Supplementary Fig. 1d).

The $c_{6A}$-overexpression strain $c_{6A}$-OE was created by transforming $c_{6A}$-KO with the constructs pDKM-000 by electroporation as described before (1) with the following specifications. A *Chlamydomonas* culture from standard growth conditions with $OD_{750}$ ~ 1 was concentrated 100-fold in TAP with 60 mM sucrose. 250 µL were then incubated with 1 µg of linearized DNA (PmeI) for 10 min on ice in an electroporation cuvette (0.4 cm gap, BioRad) prior to electroporation (BioRad Gene Pulser). Cells were recovered in 10 mL TAP with 60 mM sucrose for 16 h in a shaking incubator (2 µmol photon $m^{-2}$ $s^{-1}$, 120 rpm, 25°C) prior to plating for selection on TAP-agar plates with Paromomycin (10 µg/mL). Following selection, immunoblot analysis confirmed the expression of $c_{6A}$-FLAG in three PCR-confirmed transformants, showing a band just under 15 kDa, which corresponds well to the expected size of $c_{6A}$-FLAG without its predicted signal peptide (13.5 kDa, cut site VLA/AEA, TargetP2.0) (Supplementary Fig. 1e). One colony (number 3 in Supplementary Fig. 1e) was selected as the representative $c_{6A}$-OE strain. RNAseq analysis showed a ~14-fold increase in $c_{6A}$ transcripts compared to WT, confirming overexpression (Supplementary Fig. 1d).

The construct pDKM-000 was assembled following the molecular cloning syntax with the parts pCM0-016 (PSAD promoter) (1), pDK0-002 (*CYC4* gene with introns, this study), pCM0-094 (C-terminal triple FLAG tag) (1), and pCM0-114 (PSAD terminator) (1). The *CYC4* gene coding for $c_{6A}$ with two intronic BsaI restriction sites altered (GAGACC→GAAACC) was synthesized (Twist Bioscience).

#### PCR genotyping and whole genome sequencing

The genotype regarding the *CYC4* allele of the strains used was determined using the Phire Plant Direct PCR kit (Thermo Fisher) and following the manufacturer's instructions. Primers were used that spanned the whole $c_{6A}$ gene binding 245 bp upstream and 75 bp downstream of *CYC4* (fw: 5'-GAGACGCTACGCTGCATGT-3' & rev: 5'-ACTTCCTCGCGCTGCATA-3', annealing: 65 °C, extension: 45 s). For WT, this should lead to an amplicon of 2469 bp while for $c_{6A}$-KO a 1773 bp amplicon is expected, following the CRISPR induced deletion of 708 bp and addition of a 12 bp multiple stop codon.

For the confirmation of transformation with pDKM-000, colony PCR with the primers spanning the transgene was performed as above (fw: 5'-TCTTTCTCCATGGGCGGGCT-3', rev: 5'-CCCTCGGTCACATGTGCATCC-3', annealing: 69 °C, extension: 70 s).

For whole genome sequencing, genomic DNA from photomixotrophically grown cultures under standard conditions was obtained by phenol:chloroform extraction. The DNA was found to be of sufficient quality via spectrophotometric quantification (NanoDrop) and gel electrophoresis and was subsequently subjected to Illumina sequencing (PE-150 technology, Novogene, Cambridge, UK). After confirming the quality of the raw reads using FastQC (2), BBDuk in Geneious Prime was used to trim adapter sequences, low quality (< 30 quality value) and short reads (<10 bp). Subsequently, reads were aligned to the latest reference genome assembly for *Chlamydomonas* from JGI (CC-4532 v6.1 (3)) with the Geneious Mapper. Coverages of 97.8% (WT) and 97.4% ($c_{6A}$-KO) were reached with average read depths of 133.6x (WT) and 72.5x ($c_{6A}$-KO), and average mean MAPQ values of 202.3 (WT) and 200.3 ($c_{6A}$-KO). Integrative genomics viewer was used for coverage visualization (4).

**RNA sequencing**

Mid-exponential phase *Chlamydomonas* cultures (5 mL) were harvested by centrifugation at 1000 xg for 5 min. After disposal of the supernatant the cell concentrate was snap-frozen in liquid nitrogen. The frozen pellet was resuspended in 1 mL of TRIzol reagent (Thermo Fisher Scientific) and stored until further use at -80 °C. The TRIzol suspension was mixed with 1 mL of ethanol in preparation for RNA purification using the Direct-zol RNA Miniprep kit (Zymo Research) following the manufacturer's instructions. The optional DNase treatment and two additional wash steps were included: one wash with 0.4 mL of RNA Wash Buffer before the DNase treatment and another wash with 0.4 mL of Direct-zol RNA PreWash Buffer after the first pre-wash. The resulting concentrations and the 260/280, 260/230 ratios were subsequently quantified spectrophotometrically (NanoDrop1000, Thermo Fisher Scientific). Integrity of the extracted RNA was confirmed by gel electrophoresis.

The RNA samples were sent to Novogene for quality control, directional poly-A enriching library preparation, and Illumina sequencing (PE-150 technology). After confirming the quality of raw reads using FastQC (2), the latest genome assembly for *Chlamydomonas* from JGI (CC-4532 v6.1 (3)) was used as reference genome onto which raw reads were mapped with hisat2(5). Alignment was confirmed (between 82 and 95% aligned reads for all samples) and read assignment statistics were calculated with picard tools (6). Count data for genetic features was extracted using featureCounts (7) and the latest gff3 reference file for *Chlamydomonas* from JGI. The featureCounts table was analyzed in R, where analysis of differential expression was performed with DEseq2 (8).

**Chlorophyll quantification**

A 200 µL sample from a *Chlamydomonas* culture was mixed with 800 µL acetone, vigorously mixed for 20 s and centrifuged for 3 min at 13000 xg. The supernatant was isolated and its absorbance at 663 nm and 645 nm was measured against an 80% acetone blank. The concentration of chlorophyll a, b and total chlorophyll was calculated with the dilution factor D (here D = 5) as Chl a (µg/mL) = ($OD_{663} * 12.7 - OD_{645} * 2.69$) $*$D; Chl b (µg/mL) = ($OD_{645} * 22.9 - OD_{663} * 4.68$) $*$D; Total chl (µg/mL)= chl a + chl b (9).

**P700 redox measurements**

Simultaneous whole-cell P700 absorbance and chlorophyll fluorescence of *Chlamydomonas* (Supplementary Figure 10) were measured with a pulse amplitude-modulated spectrometer (Dual-PAM-100, Walz). Prior to measurements, cells were resuspended in TAP and the total chlorophyll concentration was adjusted to 20 µg/mL. After recovery under standard growth conditions for at least 30 min, 2 mL of the cell suspension were filtered onto glass fiber filters (WHA1822025, Whatman) and 100 µL fresh TAP were added to keep the filter moist. Measurements were performed using leaf clamps holding the filter and samples were initially incubated in darkness for 5 min before fluorescence (Intensity 12, 28 $\mu mol_{photons}$ $m^{-2}$ $s^{-1}$) and P700 measuring light (Intensity 15) were turned on.

Saturating pulses (Intensity 5, 5000 $\mu mol_{photons}$ $m^{-2}$ $s^{-1}$, 500 ms width) were applied that allowed calculation of photosynthetic parameters as before (10, 11) with P700 SP-evaluation settings adjusted to 7 ms delay and 10 ms width.

**Immunofluorescence microscopy**

Immunofluorescence staining of *Chlamydomonas* cells was carried out as previously described (12), however the methanol incubation step was left out to retain chlorophyll in the samples serving as a thylakoid marker. The antibodies used were primary mouse anti-FLAG (1:2500, B3111, Sigma) and secondary goat anti-Mouse IgG Alexa Fluor™ 488 (1:200, A11029, Invitrogen).

An inverted fluorescence microscope (Nikon Eclipse Ti) was used to investigate the stained slides. Brightfield and epifluorescence images were taken with a 20X or 100X oil immersion lens. AlexaFluor 488 signal was recorded in a YFP channel (500 nm excitation (73% intensity), yellow emission filter, 500 ms exposure) while another channel was used for chlorophyll autofluorescence (635 nm excitation (31% intensity), red emission filter, 500 ms exposure). Images were acquired as z-stacks (acquisition every 0.3 µm) and the fluorescence signals were deconvoluted using the Richardson-Lucy method and 20 iterations in the NIS-Elements Advanced Research Software (Nikon).

**Co-immunoprecipitation**

*Chlamydomonas* cultures from mid exponential growth phase were harvested by centrifugation (1000 xg, 5 min) and the pellets were resuspended in RIPA lysis buffer with freshly added PMSF (1 mM final concentration) and DNAse I (final concentration 100 U/mL). Cells were lysed by bead-beating (~100 µL of 425-600 µm beads) for 5 cycles of 1 min bead-beating and 1 min on ice. The lysate was pelleted (1 min, 2000 xg) before the supernatant was centrifuged again for 5 min at 15000 xg. The cleared lysate was then diluted with dilution buffer (10 mM Tris/Cl pH 7.5, 150 mM NaCl, 0.5 mM EDTA) and protein concentration was determined with a Bradford assay. For co-immunoprecipitation of FLAG-tagged proteins and putative interaction partners in the lysates, DYKDDDDK Fab-Trap™ Agarose beads (Chromotek) were utilized as per manufacturer's instructions with at least 4 washes and elution with 2x SDS sample buffer. For the co-immunoprecipitation of samples grown in standard conditions, a different colony than number 3 but containing the same construct (pDKM-000) was used.

**Electrochromic shift measurements of proton motive force**

Changes in the proton motive force and components, vH+ and gH+, were determined with a Dual-PAM-100 (Walz) by measuring absorbance difference signal at the 550-515 nm, known as electrochromic shift (ECS) (13). Dark interval relaxation kinetics (DIRKs) were recorded by interrupting actinic light exposure for 450 ms with darkness. Prior to measurements,

*Chlamydomonas* cells were resuspended in fresh medium and the total chlorophyll concentration was adjusted to 20 µg/mL. After recovery under standard growth conditions for at least 30 min, Ficoll 400 was added to a final concentration of 10%. Cells were dark-adapted for 10 min before measurement and measurements started with a single turnover flash (100 µs) used later for normalization of the ECS signal. For the standard growth condition induction experiment red actinic light (53 $\mu mol_{photons}$ $m^{-2}$ $s^{-1}$) was turned on and DIRKs recorded every 5 s for 30 s and every 10 s thereafter. For the DISCO Light experiment the light regime as in described in the PSII/PSI parameter measurements section was used, but instead of saturating pulses, DIRKs were recorded. At the end of the protocol, slow relaxation of the ECS signal in darkness was recorded for 1 min and used later for analysis of partitioning the pmf into ΔpH and ΔΨ (14). Analysis of kinetic data was carried out using OriginPro (Version 2024, OriginLab Corporation). The first 20 ms of the DIRK were fitted with a linear regression, the absolute slope of which corresponds to the proton flux (vH+) (15), while the inverse of the time constant of the first order exponential decay fit of a DIRK corresponds to the activity of the thylakoid conductivity (gH+) (16). ECS signals were normalized to the ECS magnitude induced by a single-turnover saturating flash of 20 µs in dark-adapted cells.

### Supplementary Figures

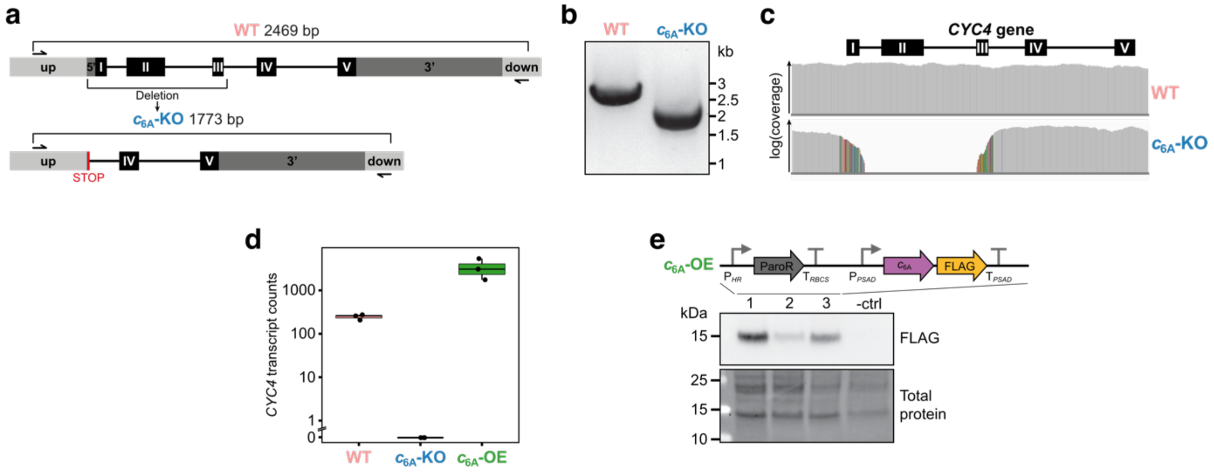


**Supplementary Figure 1: Creation and confirmation of cytochrome $c_{6A}$ mutant strains in *Chlamydomonas*. a,** The *CYC4* gene (Cre16.g670950) encoding cytochrome $c_{6A}$ was modified to create the knock-out strain $c_{6A}$-KO using CRISPR/Cpf1 which caused deletion of exons (black, roman numerals) I to III while a multiple stop codon (red, STOP) was inserted. **b,** PCR of the whole locus confirmed a deletion in $c_{6A}$-KO. **c,** Whole genome sequencing analysis of the *CYC4* locus also confirmed the CRISPR-induced deletion of exon I to III in $c_{6A}$-KO as seen from the coverage plot. The colored bars are mismatches containing the introduced multiple stop codon and bases of exon I and intron I for reads in exon III and vice versa. **d,** Counts of RNAseq reads mapped to *CYC4* comparing WT and $c_{6A}$ mutant strains when grown under continuous light confirmed $c_{6A}$-KO as knock-out and $c_{6A}$-OE as overexpression strain. **e,** Expression of C-terminally FLAG-tagged $c_{6A}$ from the GoldenGate assembled construct could be confirmed for three independent colonies but not for a negative control. Colony 3 was chosen as representative strain and named $c_{6A}$-OE for further experiments.

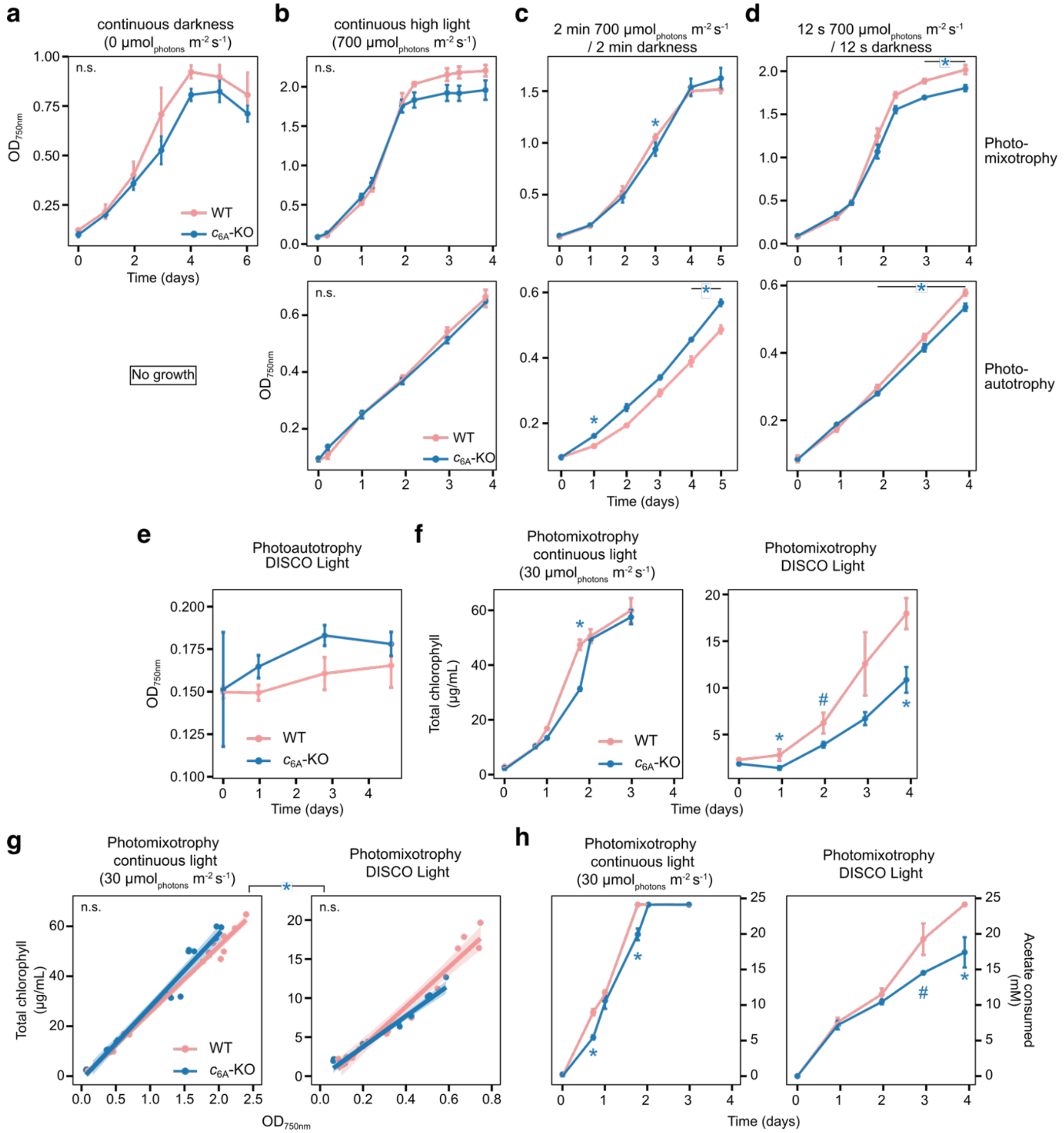


**Supplementary Figure 2: Additional growth curves, chlorophyll and acetate concentration measurements comparing WT and $c_{6A}$-KO.** Optical density ($OD_{750nm}$) growth curves of WT and $c_{6A}$-KO under various light regimes in photoautotrophy or photomixotrophy (**a**, darkness, **b**, continuous high light, **c**, 2 min high light/2 min darkness, **d**, 12 s high light/12 s darkness, all n = 3 biological replicates ±SD). **e**, Optical density ($OD_{750nm}$) growth curve of WT and $c_{6A}$ mutant strains under DISCO Light in photoautotrophy showed that none of the strains grew. DISCO Light (Darkness Interrupted by Short COnstant Light) consists of alternating 2 min darkness followed by 2 min of high light pulses (12 s high light (700 µmol$_{photons}$ m$^{-2}$ s$^{-1}$) & 12 s darkness). **f**, Comparison of total chlorophyll content per mL between WT and $c_{6A}$-KO in photomixotrophic continuous light (30 µmol$_{photons}$ m$^{-2}$ s$^{-1}$) or DISCO Light conditions (n = 3 biological replicates ±SD). **g**, Correlation between total chlorophyll concentration and $OD_{750nm}$ comparing WT and $c_{6A}$-KO in photomixotrophic continuous light (30 µmol$_{photons}$ m$^{-2}$ s$^{-1}$) or DISCO Light showed no significant differences between the strains in either condition but was different between the light treatments. **h**, Acetate consumed throughout growth in photomixotrophic continuous light (30 µmol$_{photons}$ m$^{-2}$ s$^{-1}$) or DISCO Light conditions (n = 3 biological replicates ±SD) reflected the differences observed in $OD_{750nm}$. There was 24.1

mM acetate in the medium at day 0. For the time course data (growth curves, chlorophyll per volume & consumed acetate) statistical testing was carried out with repeated measures ANOVA. Post hoc pairwise two-sided t-tests at each time point were performed if the interaction effect of time and strain on $OD_{750nm}$ in the ANOVA was significant ($p_{time:strain} < 0.05$) comparing $c_{6A}$-KO against WT ($p < 0.05$: *, $p < 0.1$: #). For the chlorophyll per OD data, three-way ANOVA with OD, strain, and light condition of the linear fits was carried out. The result of Estimated Marginal Means post-hoc testing with pairwise comparisons is indicated by asterisks ($p < 0.05$: *).

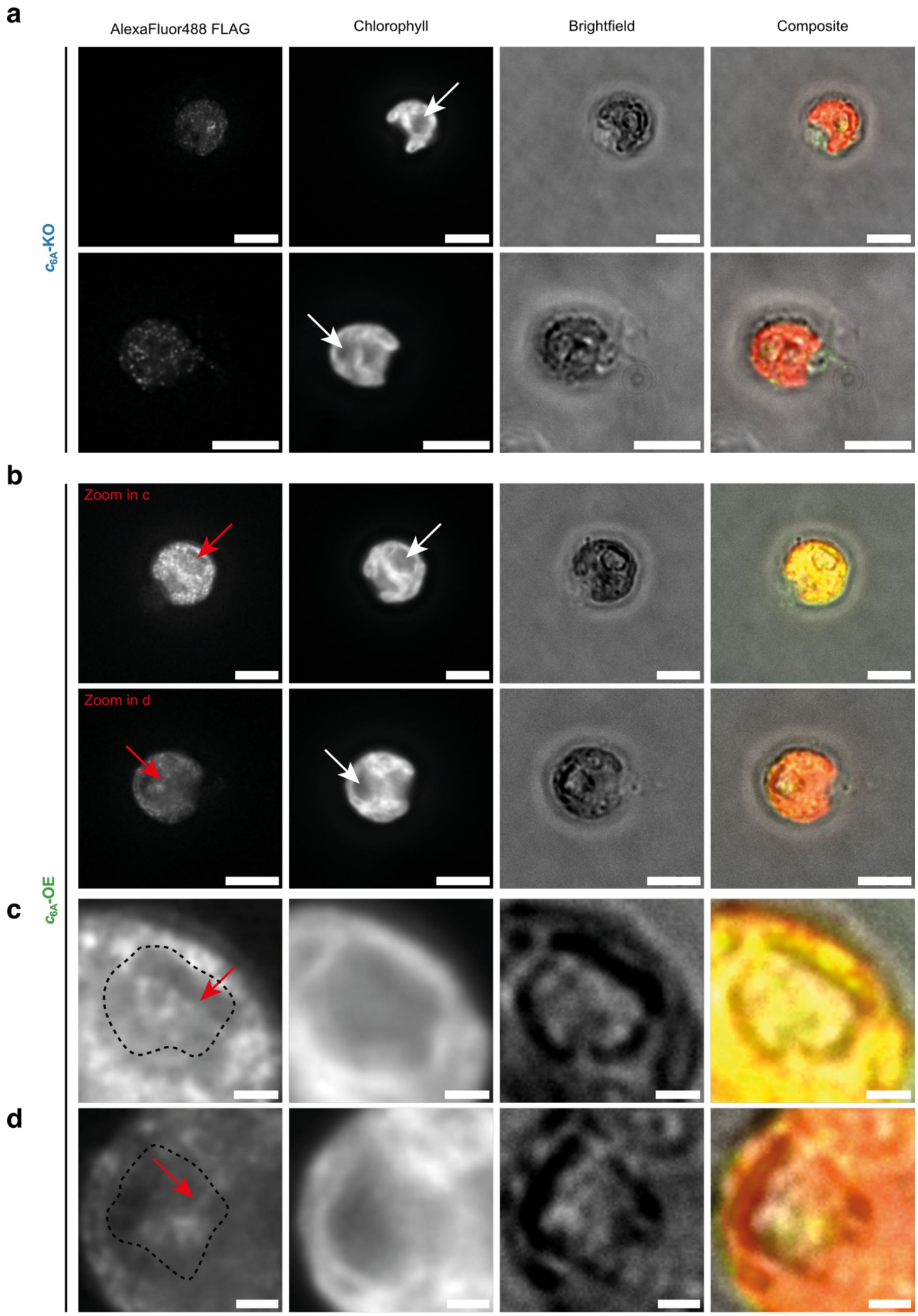


**Supplementary Figure 3: Immunofluorescence localization of $c_{6A}$-FLAG using $c_{6A}$-OE and the untransformed background strain $c_{6A}$-KO.** Two cells representing the variation of fluorescence signal are shown per strain (**a**, $c_{6A}$-KO; **b**, $c_{6A}$-OE). Samples were probed with an anti-FLAG antibody and a secondary anti-mouse antibody conjugated to AlexaFluor488 fluorescent dye. Using a 100x oil immersion lens, AlexaFluor488 (500 nm excitation, 535 nm emission filter, 500 ms exposure) and chlorophyll autofluorescence (635 nm excitation, 680 nm emission filter, 500 ms exposure) were imaged in z-stacks and a brightfield image was recorded as well (scale bar = 5 µm). Images were subjected to fluorescence signal deconvolution to improve resolution and adjusted to the same brightness and contrast values. White arrows in the chlorophyll images indicate the location of the pyrenoid. In the composite, the AlexaFluor488 and chlorophyll fluorescence are shown in green and red, respectively. **c & d**, Magnification of the images of $c_{6A}$-OE, where red arrows mark structures that resemble pyrenoid traversing thylakoid membranes (scale bar = 1 µm). The position of the starch sheath visible in the brightfield images was overlaid in broken black lines to indicate the position of the pyrenoid. Eight $c_{6A}$-OE cells were investigated more closely and all of them exhibited increased AlexaFlour488 fluorescence compared to $c_{6A}$-KO control.

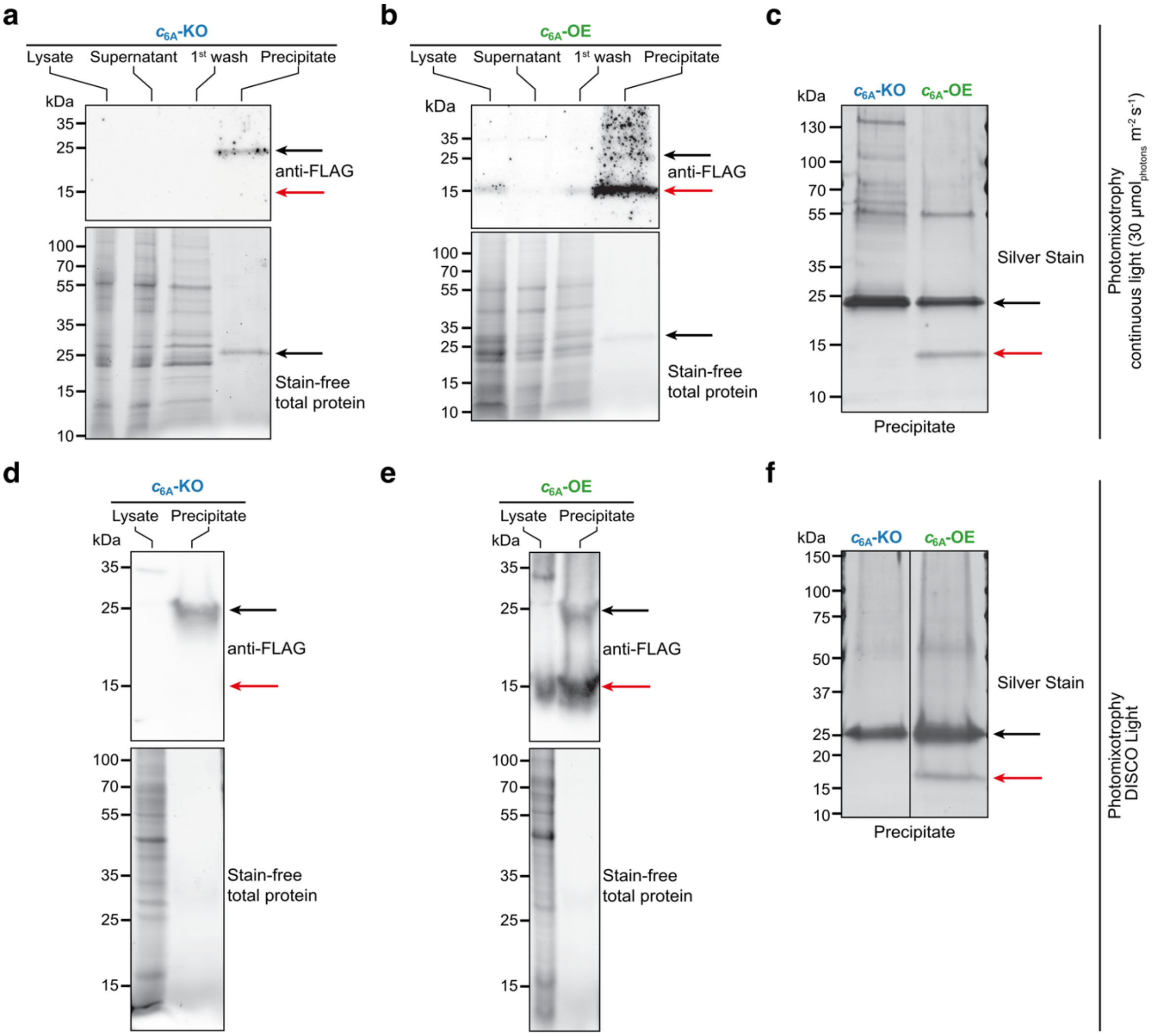


**Supplementary Figure 4: Co-immunoprecipitation of FLAG-tagged proteins does not reveal any stable interaction partners of $c_{6A}$.** **a-c,** Samples from cultures grown in photomixotrophy and continuous light (30 µmol$_{photons}$ m$^{-2}$ s$^{-1}$). **d-f,** Samples from cultures grown in photomixotrophy and DISCO Light. **a & b**, Lysate, supernatant, 1$^{st}$ wash and precipitate of **a**, $c_{6A}$-KO, and **b**, $c_{6A}$-OE, were subjected to protein gel electrophoresis (stain-free total protein) and anti-FLAG immunoblotting. $c_{6A}$-FLAG was readily detected (red arrow). **c**, An additional protein gel electrophoresis of the precipitates was silver stained. The $c_{6A}$-FLAG band is indicated by a red arrow. **d & e,** Lysate, and precipitate of **d**, $c_{6A}$-KO, and **e**, $c_{6A}$-OE, were subjected to protein gel electrophoresis (stain-free total protein) and anti-FLAG immunoblotting. **f**, An additional protein gel electrophoresis of the precipitates was silver stained. The $c_{6A}$-FLAG band is indicated by a red arrow. Black arrows indicate a 25 kDa band that appeared in all samples but was not $c_{6A}$-FLAG specific.

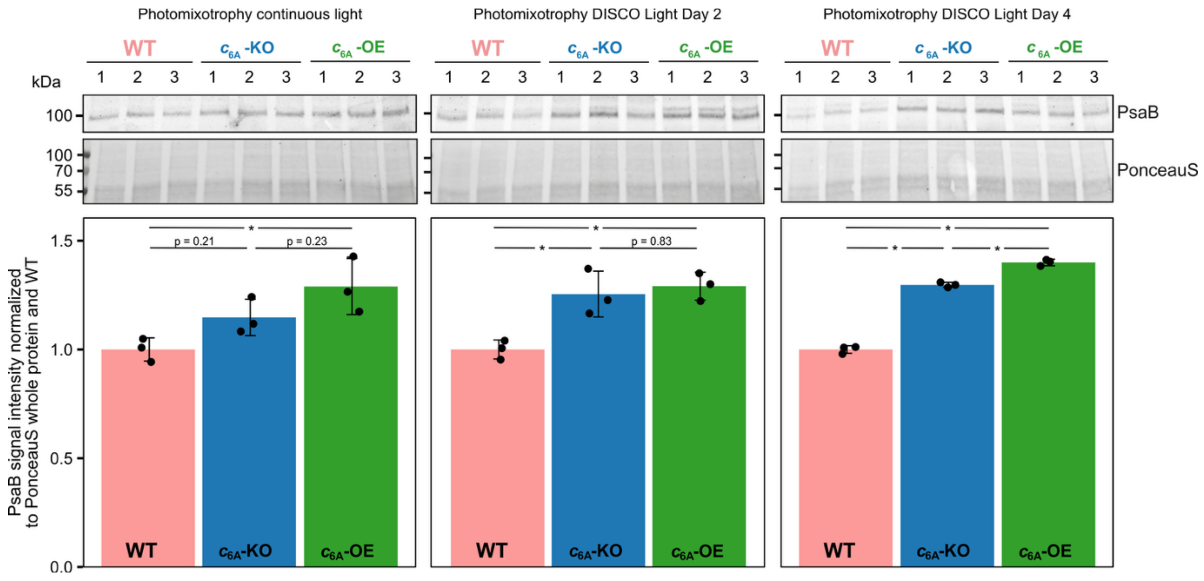


**Supplementary Figure 5: Immunoblot analyses probing for PsaB in whole cell protein samples from $c_{6A}$ mutant strains and WT grown in continuous light or DISCO Light for two and four days (n = 3 biological replicates).** For quantification signal intensity was normalized to total protein content (PonceauS stain) and WT. Asterisks indicate a p<0.05 for a two-tailed t-test comparing the samples as post-hoc test of an ANOVA.

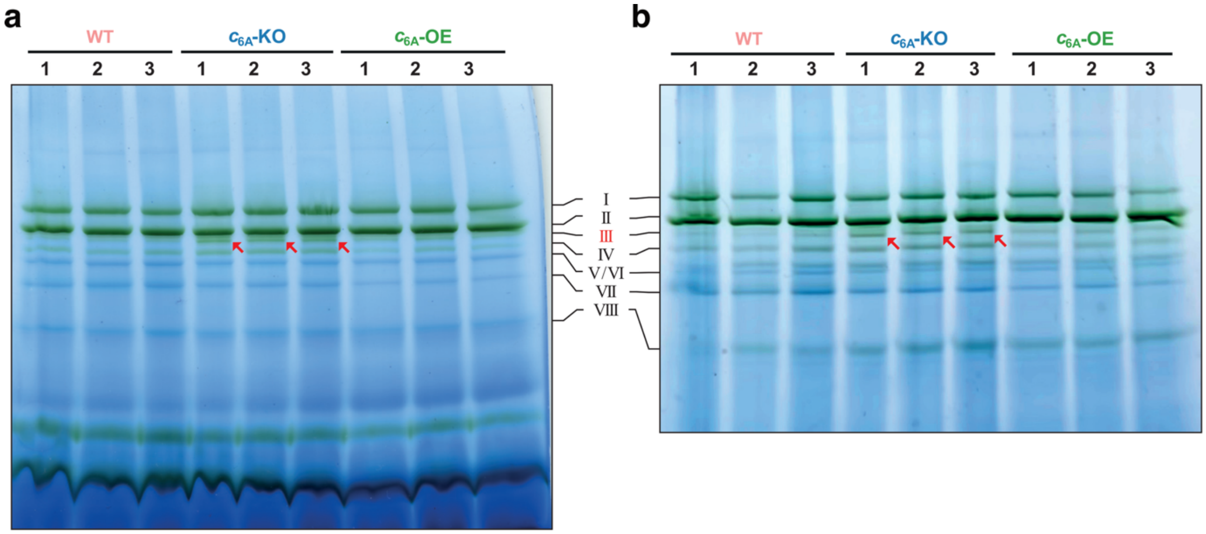


**Supplementary Figure 6: BN-PAGE analysis of $c_{6A}$ mutant strains and WT thylakoid membrane complexes isolated from cultures grown in standard conditions**. Panels **a** and **b** represent replications of the BN-PAGE with the same samples, where the gel in **b** was run longer to achieve further separation. WT3, $c_{6A}$-KO1 and $c_{6A}$-OE2 from both gels are shown in Fig. 2d and the numbering of bands (Roman numerals) corresponds to the assignment in Fig. 2d. The red arrows mark the discussed band III which was consistently more prominent in $c_{6A}$-KO samples compared to WT and $c_{6A}$-OE samples.

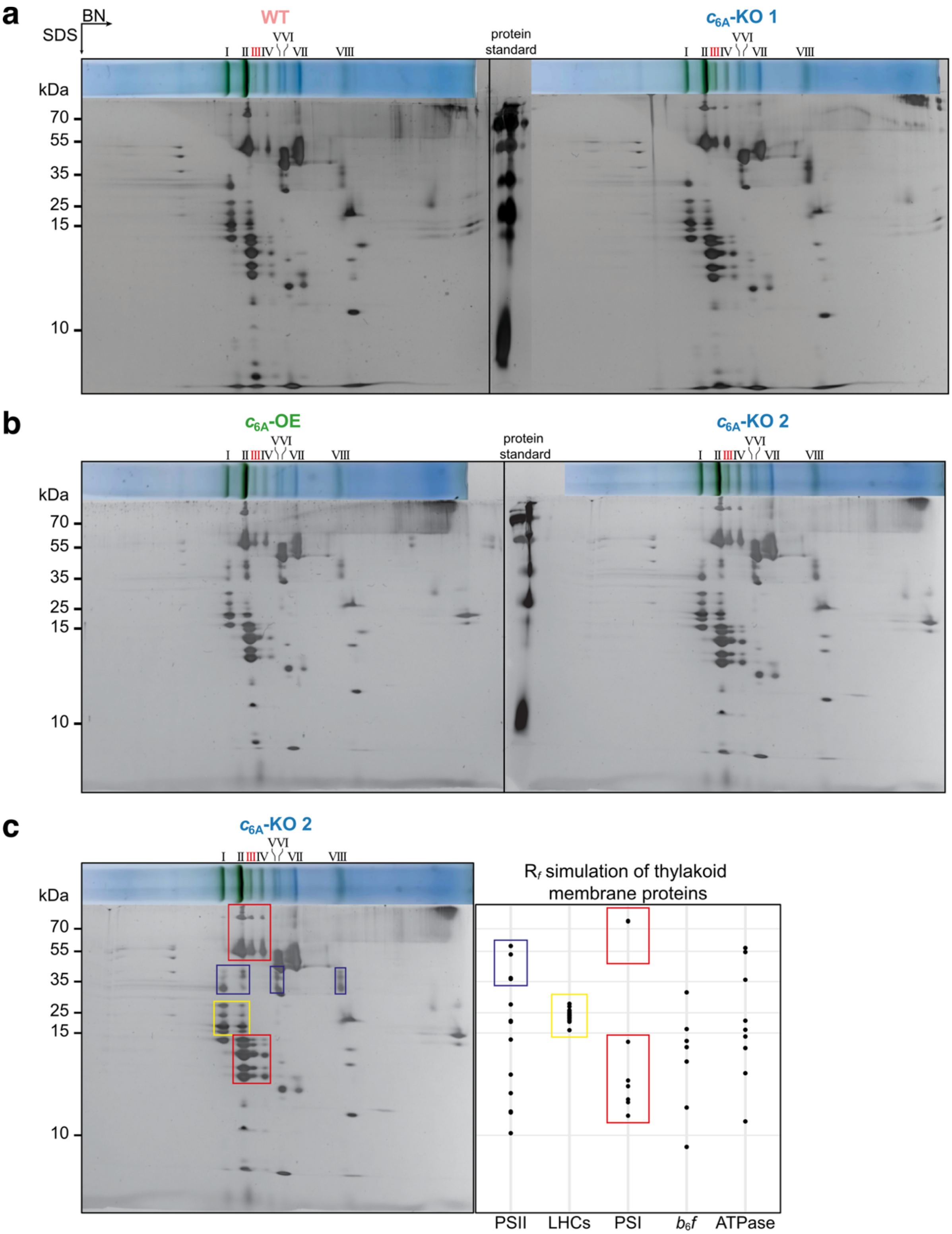


**Supplementary Figure 7: 2D-PAGE analyses of thylakoid membrane complexes from WT and $c_{6A}$ mutant strains isolated from cultures grown in standard conditions. a**, WT and $c_{6A}$-KO 1 were run on separate gels but subjected to silver stain together. Nevertheless, very similar patterns could be observed overall, and no spots were unique to either strain. **b**, Similarly, $c_{6A}$-OE and $c_{6A}$-KO 2 were run on separate gels but subjected to silver stain together. No spots were unique to either strain. **c**, Comparison of a twodimensional BN-PAGE/SDS-PAGE analysis of $c_{6A}$-KO with calculated $R_f$ values of known thylakoid membrane complex components (see also Fig. 2e). More thylakoid membrane complexes exist than those shown here but here the focus was laid on complexes involved in photosynthesis and especially photosystems and antennae (LHCs), as the band of interest (III) was green. The presence of bands at 55 kDa and above 70 kDa for bands III and IV is not reflected in the $R_f$ simulation, where only two spots (PsaA & PsaB) were expected above 70 kDa. However, previous mass spectrometric analyses have also found PsaA at two positions of different high molecular weight, too (17).

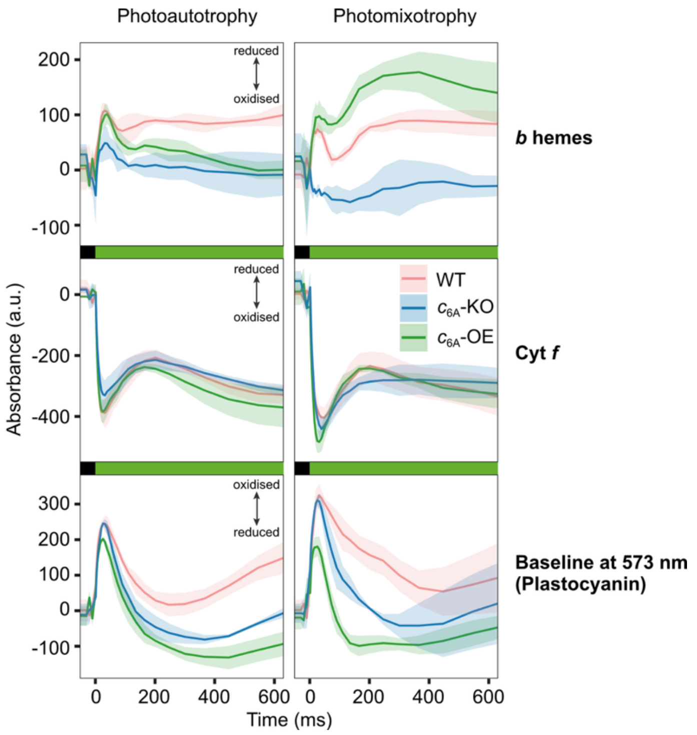


**Supplementary Figure 8: Redox changes of *b* hemes, cytochrome *f*, and plastocyanin in $c_{6A}$ mutant strains and WT.** Photoautotrophic or photomixotrophic exponential growth phase cultures were exposed to green light (500 µmol$_{photons}$ m$^{-2}$ s$^{-1}$, green bar) in a JTS spectrometer (n = 3 biological replicates ±SD). A plastocyanin signal is recorded as part of the baseline measurement at 573 nm, although its peak absorption is at 597 nm. The plastocyanin signal shown here is thus not the focus of the measurements. However, differences between the strains could be observed.

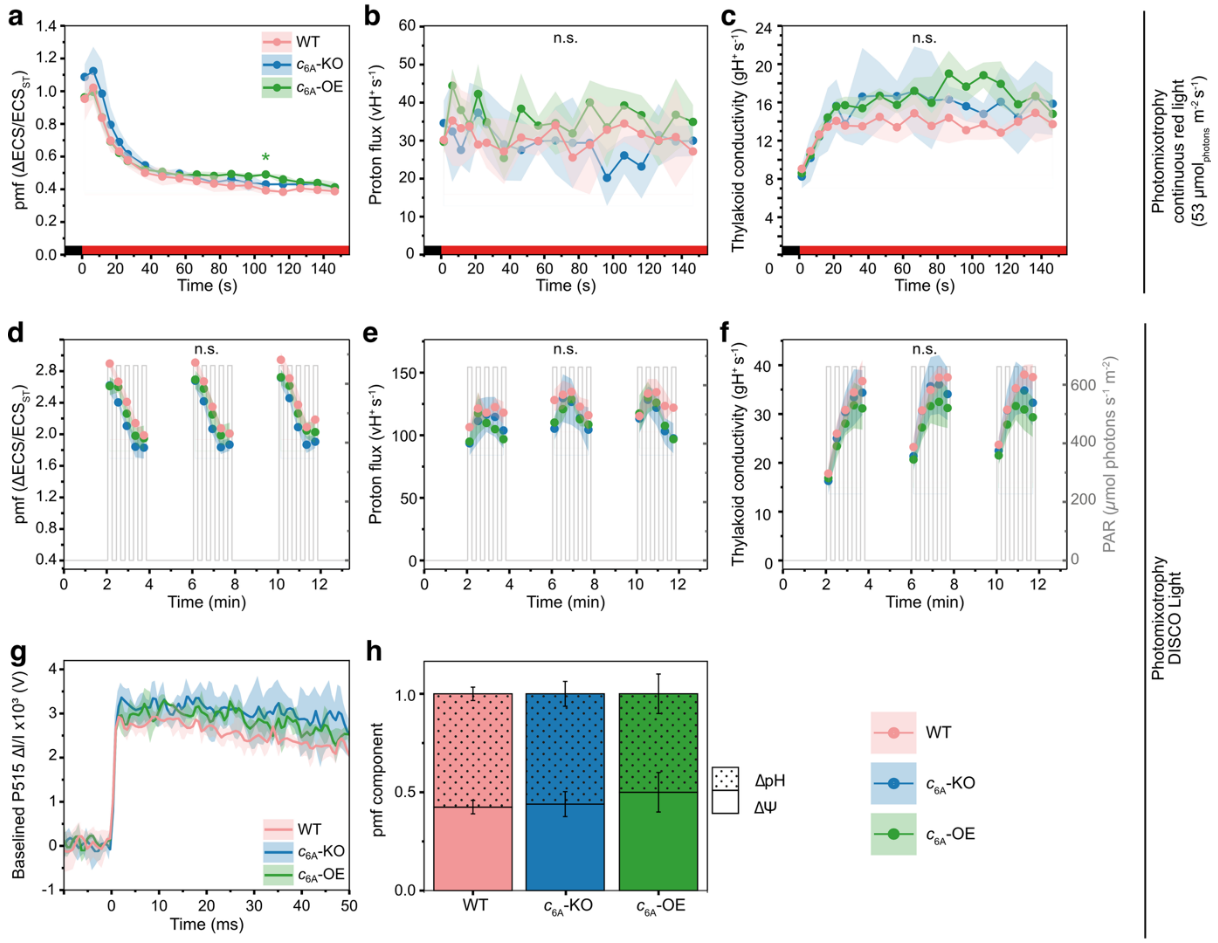


**Supplementary Figure 9: Electrochromic shift (ECS) measurements of $c_{6A}$ mutant strains and WT did not show substantial differences in either continuous or DISCO Light. a**, Proton motive force pmf, **b**, proton flux vH$^+$, and **c**, thylakoid conductivity gH$^+$ of $c_{6A}$ mutant strains and WT grown in photomixotrophic continuous light (30 µmol$_{photons}$ m$^{-2}$ s$^{-1}$) conditions and exposed to red light (53 µmol$_{photons}$ m$^{-2}$ s$^{-1}$, red bar, n = 3 biological replicates ±SD). The pmf data is normalized to the ECS signal change caused by a single turnover flash before the continuous illumination. **d**, Proton motive force pmf, **e**, proton flux vH$^+$, and **f**, thylakoid conductivity gH$^+$ of $c_{6A}$ mutant strains and WT grown in photomixotrophic continuous light (30 µmol$_{photons}$ m$^{-2}$ s$^{-1}$) conditions and exposed to DISCO Light (secondary y-axis, n = 3 biological replicates ±SD). The pmf data is normalized to the ECS signal change caused by a single turnover flash before the continuous illumination (see **g**). Non-significance is indicated by "n.s." and the green colored asterisk marks a significant difference ($p < 0.05$) of $c_{6A}$-OE to $c_{6A}$-KO as determined by repeated measures ANOVA and pairwise two-sided t-tests as post-hoc test if the ANOVA $p_{time:strain}$ value was below 0.05. **g**, The ECS response to a single turnover flash (applied at t = 0 ms) was recorded before the start of the DISCO Light protocol. **h**, The partition of pmf into ΔpH and ΔΨ was determined at the end of 12 minutes of the DISCO light protocol.

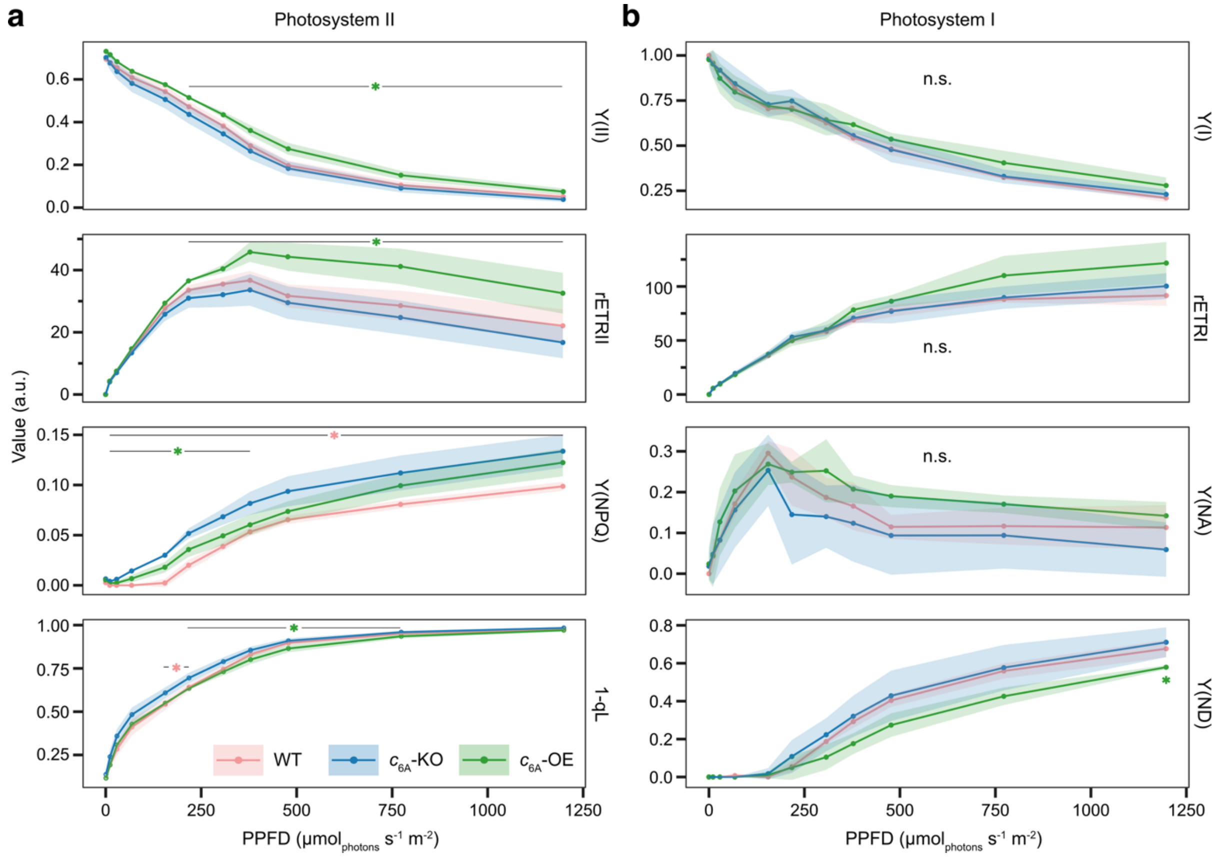


**Supplementary Figure 10: Light response curves of $c_{6A}$ mutant strains and WT. a**, Cultures grown in standard conditions were measured for their photosystem II quantum yield Y(II), relative electron transfer rate rETRII, regulated non-photochemical quenching yield Y(NPQ), and QA reduction state 1-qL (a proxy for the PQ pool redox state) using a Dual-PAM-100 (n = 3 biological replicates ±SD). **b**, Simultaneously, the samples were also assessed for their photosystem I quantum yield Y(I), rETRI, acceptor side limitation Y(NA) and donor-side limitation Y(ND) (n = 3 biological replicates ±SD). The relative electron transfer rates (rETR) for PSII and PSI were calculated as photosystem quantum yield x photosynthetic photon flux density (PPFD). Non-significance is indicated by "n.s." and colored asterisks mark a significant difference ($p < 0.05$) of WT (pink) or $c_{6A}$-OE (green) to $c_{6A}$-KO as determined by repeated measures ANOVA and pairwise two-sided t-tests as post-hoc test if the ANOVA $p_{PAR:strain}$ value was below 0.05.

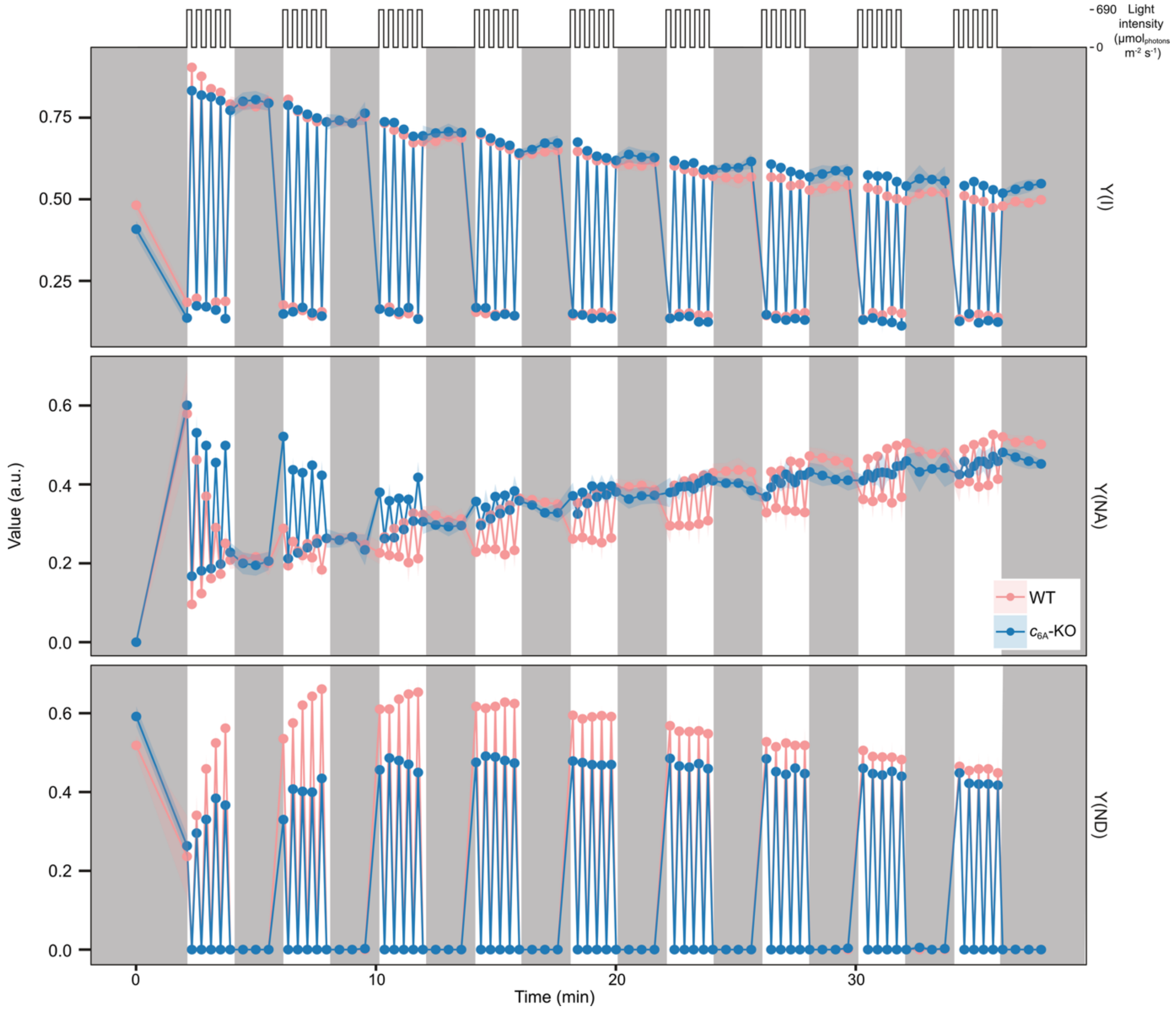


**Supplementary Figure 11: Measurements of photosystem I parameters of $c_{6A}$-KO and WT under DISCO Light conditions**. Photomixotrophic exponential growth phase cultures (adjusted to 20 µg/mL total chlorophyll) were applied to a glass fiber filter and exposed to a DISCO Light regime in a Dual-PAM-100 spectrometer where grey boxes represent darkness and white boxes represent 2 min of 12 s on/off as depicted above the Y(I) panel (N=3 ±SD). The effect of saturating pulses on P700 differential absorbance was used to calculate photosystem I quantum yield Y(I), acceptor-side limitation Y(NA) and donor-side limitation Y(ND).

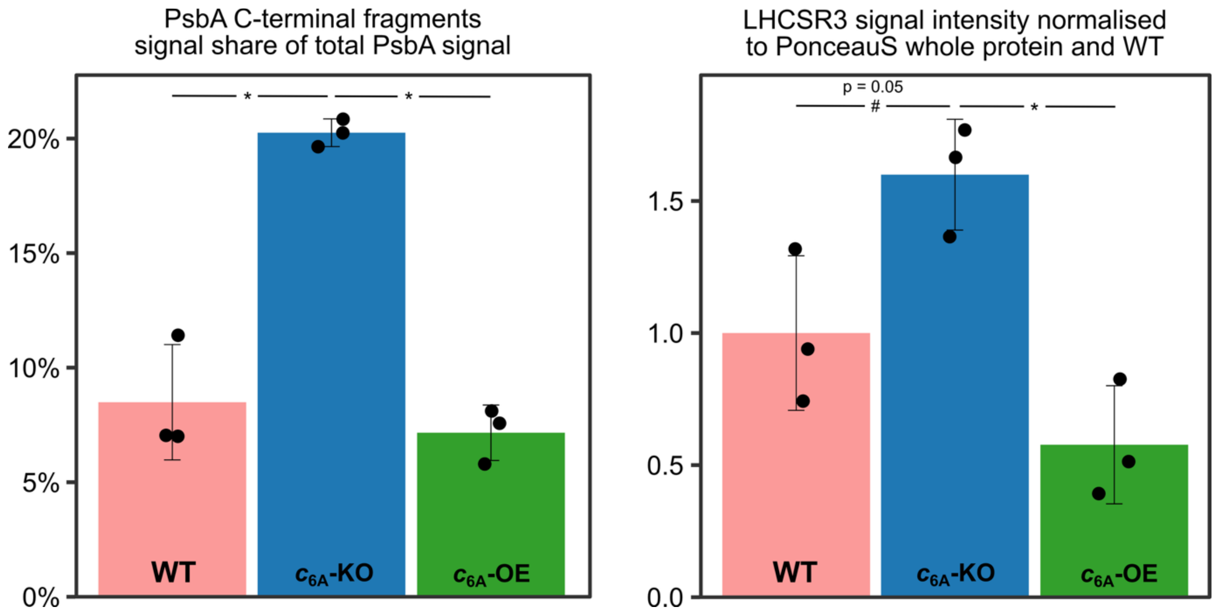


**Supplementary Figure 12: Quantification of western blot band intensities.** For PsbA C-terminal fragments: relative to total PsbA signal, for LHCSR3: relative to total protein (PonceauS). Asterisks indicate a significant difference ($p < 0.05$) as determined by one-way ANOVA and post-hoc pairwise two-sided t-test ($n = 3$ biological replicates ±SD).

**Supplementary Table 1: Summary of experimental findings ordered by figure and method.**

| Figure | Method | Finding |
|---|---|---|
| **1** | Growth curves | $c_{6A}$-dependent growth phenotype only under DISCO Light (absence →less growth) |
| **2** | 77K fluorescence | $c_{6A}$-dependent light harvesting balance phenotype under normal conditions in photomixotrophy (absence →lower PSI/PSII fluorescence ratio) |
| **2** | 77K fluorescence | Ability for state transitions is still present but $c_{6A}$-KO always had lower PSI/PSII fluorescence ratio in either state in photomixotrophy |
| **2** | BN-PAGE & 2D-PAGE | $c_{6A}$-dependent difference in accumulation of a likely LHC-less PSI complex (absence →more prominent complex) |
| **3** | OJIP & DUAL-PAM | More reduced PQ pool in the absence of $c_{6A}$ |
| **3** | JTS | Light-induced oxidation of *b* hemes (esp. in photomixotrophy) in the absence of $c_{6A}$, which is explainable by the effect on the PQ pool |
| **3** | DUAL-PAM | Under DISCO Light, absence of $c_{6A}$ leads to decreased PSII quantum yield |
| **4** | DUAL-PAM | Higher values of stress indicators Y(NPQ) & Y(NO) in DISCO Light in the absence of $c_{6A}$ |
| **4** | Western blot | Higher levels of stress indicators LHCSR3 & PsbA fragments in DISCO Light in the absence of $c_{6A}$ |
| **Supplementary 1** | PCR, whole genome sequencing, RNAseq, Western blot | $c_{6A}$-KO and $c_{6A}$-OE strains were confirmed to be knock-out and overexpression lines, respectively |
| **Supplementary 2** | Growth curves, chlorophyll & acetate quantification | No $c_{6A}$-dependent growth phenotype under darkness or other fluctuating light while acetate and chlorophyll contents reflect DISCO Light phenotype |
| **Supplementary 3** | Immunofluorescence microscopy | $c_{6A}$ was likely thylakoid luminal localized |
| **Supplementary 4** | Co-IP | $c_{6A}$ did not form stable complexes with other proteins |
| **Supplementary 5** | Western blot | PsaB is not reduced in $c_{6A}$-KO compared to WT |
| **Supplementary 6** | BN-PAGE | Detection of likely LHC-less PSI complex was consistently increased in $c_{6A}$-KO compared to WT and $c_{6A}$-OE |
| **Supplementary 7** | 2D-PAGE | Composition of $c_{6A}$-dependently over-accumulated complex suggested it is LHC-less PSI complex |
| **Supplementary 8** | JTS | $c_{6A}$-dependent changes in *b* hemes redox kinetics were not reflected in cytochrome *f* and plastocyanin redox kinetics |
| **Supplementary 9** | ECS | No $c_{6A}$-dependent difference in pmf, $gH^+$, $vH^+$ in DISCO Light |
| **Supplementary 10** | DUAL-PAM | $c_{6A}$-dependent difference in NPQ (absence →slightly elevated NPQ) and PQ pool redox state (absence →more reduced PQ pool) in continuous light |
| **Supplementary 11** | DUAL-PAM | Small $c_{6A}$-dependent differences in PSI under DISCO Light (→ consistent with more reduced PQ pool) |
| **Supplementary 12** | Western blot | Quantification confirms higher levels of stress indicators LHCSR3 & PsbA fragments in DISCO Light in the absence of $c_{6A}$ |

### SI References


1. P. Crozet, *et al.*, Birth of a Photosynthetic Chassis: A MoClo Toolkit Enabling Synthetic Biology in the Microalga *Chlamydomonas reinhardtii*. *ACS Synth. Biol.* **7**, 2074–2086 (2018).

2. S. Andrews, FastQC: A Quality Control Tool for High Throughput Sequence Data. (2010). Available at: http://www.bioinformatics.babraham.ac.uk/projects/fastqc/.

3. R. J. Craig, *et al.*, The Chlamydomonas Genome Project, version 6: Reference assemblies for mating-type plus and minus strains reveal extensive structural mutation in the laboratory. *Plant Cell* **35**, 644–672 (2023).

4. J. T. Robinson, *et al.*, Integrative genomics viewer. *Nat. Biotechnol.* **29**, 24–26 (2011).

5. D. Kim, J. M. Paggi, C. Park, C. Bennett, S. L. Salzberg, Graph-based genome alignment and genotyping with HISAT2 and HISAT-genotype. *Nat. Biotechnol.* **37**, 907–915 (2019). Available at: http://dx.doi.org/10.1038/s41587-019-0201-4.

6. Broad Institute, Picard Tools. Available at: http://broadinstitute.github.io/picard.

7. Y. Liao, G. K. Smyth, W. Shi, FeatureCounts: An efficient general purpose program for assigning sequence reads to genomic features. *Bioinformatics* **30**, 923–930 (2014).

8. M. I. Love, W. Huber, S. Anders, Moderated estimation of fold change and dispersion for RNA-seq data with DESeq2. *Genome Biol.* **15**, 1–21 (2014).

9. D. I. Arnon, COPPER ENZYMES IN ISOLATED CHLOROPLASTS. POLYPHENOLOXIDASE IN *BETA VULGARIS*. *Plant Physiol.* **24**, 1–15 (1949).

10. C. Klughammer, U. Schreiber, Saturation Pulse method for assessment of energy conversion in PS I. *PAM Appl. Notes* 11–14 (2008).

11. C. Klughammer, U. Schreiber, Complementary PS II quantum yields calculated from simple fluorescence parameters measured by PAM fluorometry and the Saturation Pulse method. *PAM Appl. Notes* **1**, 27–35 (2008).

12. T. Yamano, H. Fukuzawa, Indirect Immunofluorescence Assay in Chlamydomonas reinhardtii. *BIO-Protoc.* **6**, 1–6 (2016).

13. B. Bailleul, P. Cardol, C. Breyton, G. Finazzi, Electrochromism: A useful probe to study algal photosynthesis. *Photosynth. Res.* **106**, 179–189 (2010).

14. J. A. Cruz, C. A. Sacksteder, A. Kanazawa, D. M. Kramer, Contribution of electric field ($\Delta\psi$) to steady-state transthylakoid proton motive force (pmf) in vitro and in vivo. Control of pmf parsing into $\Delta\psi$ and $\Delta$pH by ionic strength. *Biochemistry* **40**, 1226–1237 (2001).

15. P. Joliot, A. Joliot, Cyclic electron transfer in plant leaf. *Proc. Natl. Acad. Sci. U. S. A.* **99**, 10209–10214 (2002).

16. A. Kanazawa, D. M. Kramer, In vivo modulation of nonphotochemical exciton quenching (NPQ) by regulation of the chloroplast ATP synthase. *Proc. Natl. Acad. Sci. U. S. A.* **99**, 12789–12794 (2002).

17. S. Rexroth, J. M. W. Meyer zu Tittingdorf, F. Krause, N. A. Dencher, H. Seelert, Thylakoid membrane at altered metabolic state: Challenging the forgotten realms of the proteome. *Electrophoresis* **24**, 2814–2823 (2003).
